# Parasite-induced ER-PVM contacts promote CERT-dependent ceramide delivery to the apicomplexan vacuole

**DOI:** 10.64898/2026.11.13.763891

**Authors:** Vaisak Mohan, Ilaksha Pahwa, Veeda Narahari, Ajmal P Mustafa, Purvit Bhatia, Sahoo Parnika, Usha Krishna, Bharathi Bhogapurapu, Manish Kumar, Kota Arun Kumar, Kamalakannan Vijayan

## Abstract

Apicomplexan parasites rely on host lipids for replication, yet how host transport machinery is coupled to the parasitophorous vacuole membrane (PVM) remains unresolved. Here, we uncover a host-parasite membrane contact site mechanism driving ceramide delivery to the vacuole. Host ceramide transfer protein (CERT) targets the PVM via its PH domain, cooperating with host VAP-B to promote sphingolipid accumulation and support *Toxoplasma* and *Plasmodium* development. By combining VAP-B proximity labeling with FFAT motif predictions, we identified TgVPS13 as a parasite-encoded FFAT effector that physically engages host VAP-B at the host-parasite interface. Conditional depletion of TgVPS13 markedly reduces vacuolar sphingolipid enrichment, partially impairs host CERT recruitment at the PVM, disrupts intravacuolar parasite organization, and severely restricts replication. Together, these findings establish a model in which TgVPS13-VAP-B tethering creates ER-PVM proximity, organizing a functional contact site that optimizes CERT-mediated ceramide transfer for efficient host lipid acquisition.

## Main

Apicomplexan parasites, such as *Toxoplasma* and *Plasmodium* species, rely heavily on scavenging host cell lipids to construct their parasitophorous vacuole membrane (PVM) and sustain rapid intracellular replication^1–3^. Among these host-derived lipids, sphingolipids play important roles in parasite fitness, and infection can remodel host sphingolipid metabolism and increase sphingolipid availability^4,5^. Because primary ceramide synthesis is compartmentalized at the host endoplasmic reticulum (ER), a fundamental question is how parasites access ER-derived ceramide and direct its delivery to the PVM^6^.

Genetic and functional studies have implicated the host ceramide transfer protein CERT in apicomplexan development^7,8^. A genome-wide CRISPR-Cas9 screen in liver-stage *Plasmodium* identified CERT among host factors required for optimal parasite development^7^. More recently, host sphingolipids were shown to support *P. berghei* liver-stage development, with exogenous ceramide promoting parasite growth and CERT localizing to the parasite vacuole^8^. However, the spatial mechanism that allows a PVM-associated lipid transfer protein to efficiently access ceramide from the host ER remains unclear^9,10^.

Recent studies established that *T. gondii* actively recruits host ER machinery to the parasite vacuole through parasite-encoded effectors. TgROP1 functions as a parasite-encoded tether that binds the MSP domains of host VAPA and is required for ER-*Toxoplasma* membrane contact sites^11,12^. Independently, TgVIP1 was identified as a parasite protein containing an FFAT-like motif that interacts with VAPB^12^, while VAPA, VAPB, and MOSPD2 contribute to ER-PV interactions^12,13^. These studies establish ER-PVM contacts as an important feature of the parasite-host interface, but whether these contacts functionally couple ER lipid availability to dedicated lipid transfer machinery remains unresolved.

Here, we investigated the ER-PVM contact sites as functional platforms for host sphingolipid delivery^14^. We show that host-derived sphingolipids are acquired by both *Toxplasma* and *Plasmodium*, and that CERT targets the PVM through its PH domain and is required for parasite development and survival, *in vitro* and *in vivo*. To identify parasite-side machinery associated with this lipid-transfer interface, we performed unbiased *in situ* proximity labelling using miniTurbo mass spectrometry^15^ and identified TgVPS13 as a parasite-encoded, FFAT-containing protein^16^ that associates with the host ER protein VAP-B. Conditional degradation of TgVPS13 markedly reduces vacuolar sphingolipid enrichment, partially disrupts host CERT recruitment at the vacuolar interface, causes disorganization of intravacuolar parasite architecture, and severely impairs parasite replication.

Our findings support a model in which TgVPS13-VAP-B-mediated tethering brings the host ER into close proximity with the parasitophorous vacuole, establishing a spatial environment that optimizes CERT-mediated transfer of ER-derived ceramide. By linking parasite-encoded membrane contact site machinery directly to a defined host lipid transfer pathway, this work reveals how apicomplexan parasites exploit host organelle contact machinery to efficiently acquire essential host lipids.

### Host ER closely associates with the apicomplexan vacuole and host-derived sphingolipids accumulate at the PVM

To examine the spatial relationship between host ER membranes and the parasite compartment, we performed confocal imaging of intracellular *T. gondii* and liver-stage *P. yoelii*. In both apicomplexan models, host ER networks closely surrounded the parasite vacuole, displaying extensive signal around the vacuolar boundary throughout intracellular development (**Fig. S1A**). These observations align with the established spatial association between the host ER and the parasite vacuole across apicomplexans.

We next asked whether host-derived sphingolipids access the parasite vacuole. To distinguish newly supplied lipids from pre-existing host lipid pools, we deployed complementary labelling strategies in *T. gondii* (**Fig. S1B**). Post-invasion addition of BODIPY-ceramide at 4 h post-infection resulted in pronounced accumulation of ceramide-derived fluorescence at the parasite vacuole by 36 h post-infection (**Fig. S1B**). Spatial line-scan analysis confirmed strong signal enrichment at the vacuolar boundary (**Fig. S1B**, arrowheads).

To determine whether parasites tap into pre-existing host sphingolipid pools, host cells were pre-labelled with BODIPY-ceramide for 24 h and thoroughly washed prior to parasite inoculation (**Fig. S1B**). Despite removal of extracellular probe before invasion, ceramide-associated fluorescence clearly concentrated at the parasite vacuole following intracellular growth, indicating that parasites can access pre-existing host sphingolipid pools (**Fig. S1B**).

Similarly, host cells pre-labelled with fluorescent sphingosine exhibited strong fluorescence enrichment at the parasitophorous vacuole membrane (**Fig. S1B**). Because the acylation of sphingosine into ceramide occurs predominantly via host ER-resident ceramide synthases, its selective concentration at the vacuole indicates that host ER-processed lipid products are systematically trafficked across the ER-PVM interface.

Together, these findings establish that host-derived sphingolipids accumulate at the apicomplexan vacuole, with parasites accessing both pre-existing and metabolically processed host lipid pools. The close apposition of host ER networks with the parasite vacuole points to a spatial route for lipid transfer, raising the question of whether specific ER-PVM contact site machinery mediates this delivery.

### Host ceramide transfer protein CERT is recruited to the parasitophorous vacuole in apicomplexan parasites

The accumulation of host-derived sphingolipids at the parasite vacuole and the close association of the host ER with the PVM prompted us to investigate whether a host lipid transfer protein is positioned at this interface. Genome-wide CRISPR-Cas9 screening previously identified the ceramide transfer protein CERT (*COL4A3BP*) as a host factor required for *Plasmodium* liver-stage infection^7^, suggesting that CERT may contribute to host ceramide trafficking to the parasite compartment. We therefore examined host CERT localization during apicomplexan infection. Endogenous CERT antibody staining demonstrated distinct rim-like enrichment surrounding GFP-expressing *T. gondii* vacuoles at both 30 and 44 hpi (**Fig. 1A**, insets). To independently confirm this localization, we engineered host cells to express mCherry-tagged CERT and subjected them to *T. gondii* infection. Ectopically expressed CERT-mCherry accumulated around the parasite vacuole as early as 8 hpi and remained associated with the vacuolar interface at 44 hpi, recapitulating the pattern observed for endogenous CERT (**Fig. 1B**).

**Figure 1.**
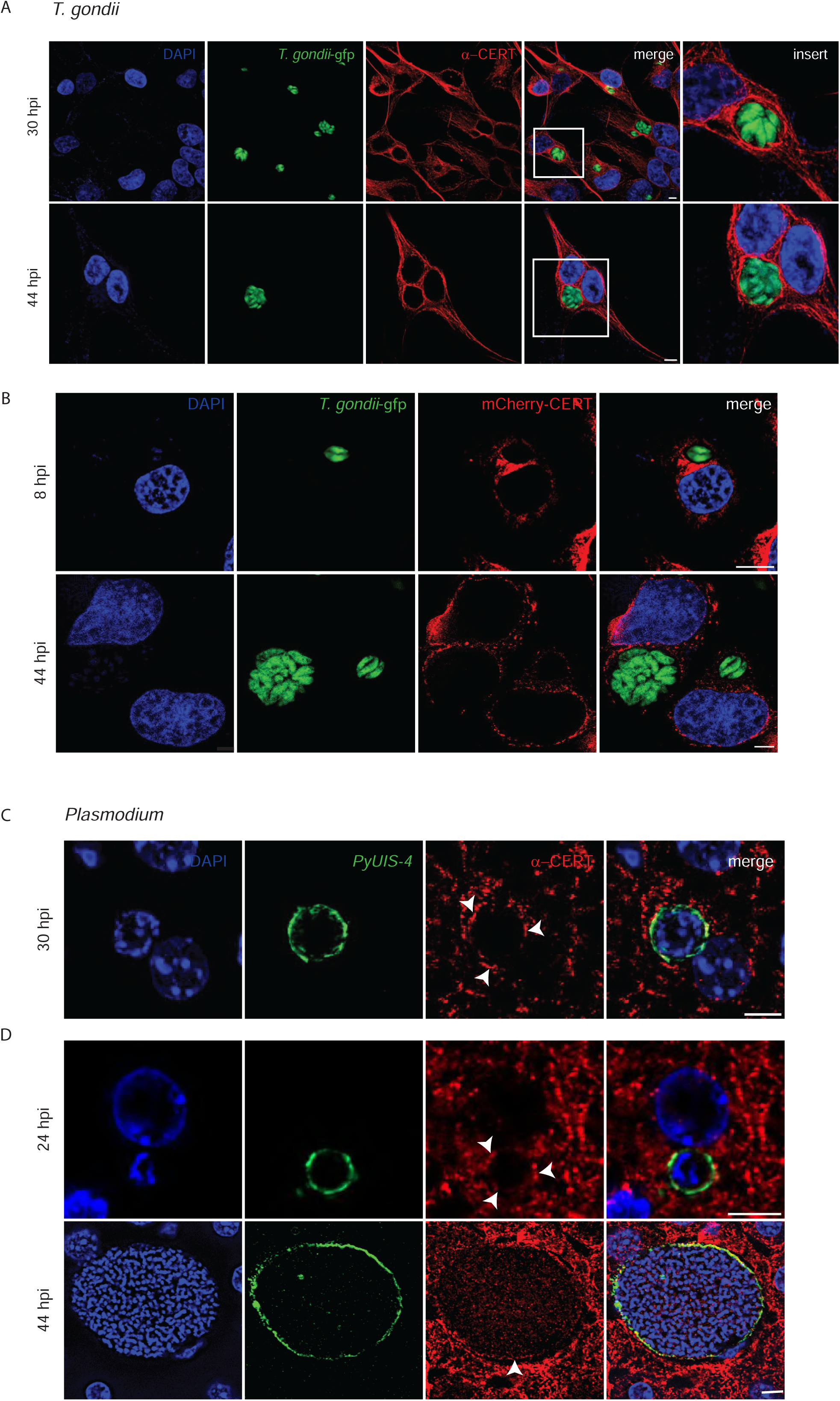
Host ceramide transfer protein CERT is recruited to the parasitophorous vacuole across apicomplexan species. **(A)** Confocal immunofluorescence images showing endogenous CERT localization in host cells infected with GFP-expressing *T. gondii* at 30 and 44 hpi. Insets highlight rim-like CERT enrichment around the parasite vacuole. **(B)** Fluorescence images of host cells expressing CERT-mCherry infected with *T. gondii* at 8 and 44 hpi, independently confirming the association of ectopically expressed CERT with the parasite vacuole. **(C)** Confocal immunofluorescence images showing endogenous CERT localization in Hepa1-6 cells infected with liver-stage *P. yoelii* at 30 hpi. Parasite vacuoles are co-stained with the PVM marker *Py*UIS-4. Arrowheads indicate CERT enrichment at the PVM. **(D)** Immunofluorescence images showing endogenous CERT localization at the *P. yoelii* liver-stage vacuole in infected mouse liver at early (24 hpi) and late (44 hpi) stages. Arrowheads indicate CERT enrichment at the parasite vacuole. Scale bars, 5 μm.

We next examined CERT localization during liver-stage *P. yoelii* infection. In infected Hepa1-6 cells, immunofluorescence microscopy revealed prominent enrichment of endogenous CERT at the PVM as demonstrated previously^8^. At 30 hpi, the CERT signal closely coincided with the vacuolar boundary marked by PyUIS-4 (**Fig. 1C**, arrowheads). We then asked whether this association was also evident during *P. yoelii* infection *in vivo*. In liver sections from infected mice, endogenous CERT was enriched at the parasite vacuole at both early (24 hpi) and late (44 hpi) stages of liver-stage development (**Fig. 1D**, arrowheads). Thus, CERT association with the *Plasmodium* liver-stage vacuole is observed both in hepatocytes *in vitro* and during infection *in vivo*, and is maintained across distinct stages of parasite development.

Together, these results demonstrate that host CERT is enriched at the parasite vacuole in both *T. gondii* and *Plasmodium*. The conserved localization of CERT at the vacuolar interface places host ceramide transfer machinery at a site of close host-parasite membrane contact where host-derived sphingolipids accumulate, suggesting that CERT may contribute to lipid trafficking at the parasite vacuole.

### CERT-dependent ceramide transport supports apicomplexan development and *in vivo* parasite growth

Having established that CERT is recruited to the parasite vacuole in both *Plasmodium* and *Toxoplasma*, we next investigated the functional requirement for CERT-dependent ceramide transport during intracellular development. Host-side genome-wide screens and recent findings^7^ identified CERT as a host dependency in liver-stage *P. berghei^8^*. To confirm and extend these findings in *P. yoelii*, we evaluated parasite development following genetic knockout of CERT (CERT K/o) (**Fig. S2A**) and pharmacological inhibition using HPA-12. While initial parasite entry at 90 min post-infection remained unaffected, CERT K/o significantly reduced overall infection at 48 h post-infection (**Fig. 2A**). Consistent with impaired intracellular development, CERT depletion markedly decreased exoerythrocytic form (EEF) size at 48 hpi (**Fig. 2B**). Pharmacological inhibition with HPA-12 recapitulated these phenotypes, with no significant effect on infection at 90 min but a significant reduction in infection rate (Fig. 3C) and EEF growth at 48 hpi (**Fig. 2D**).

**Figure 2.**
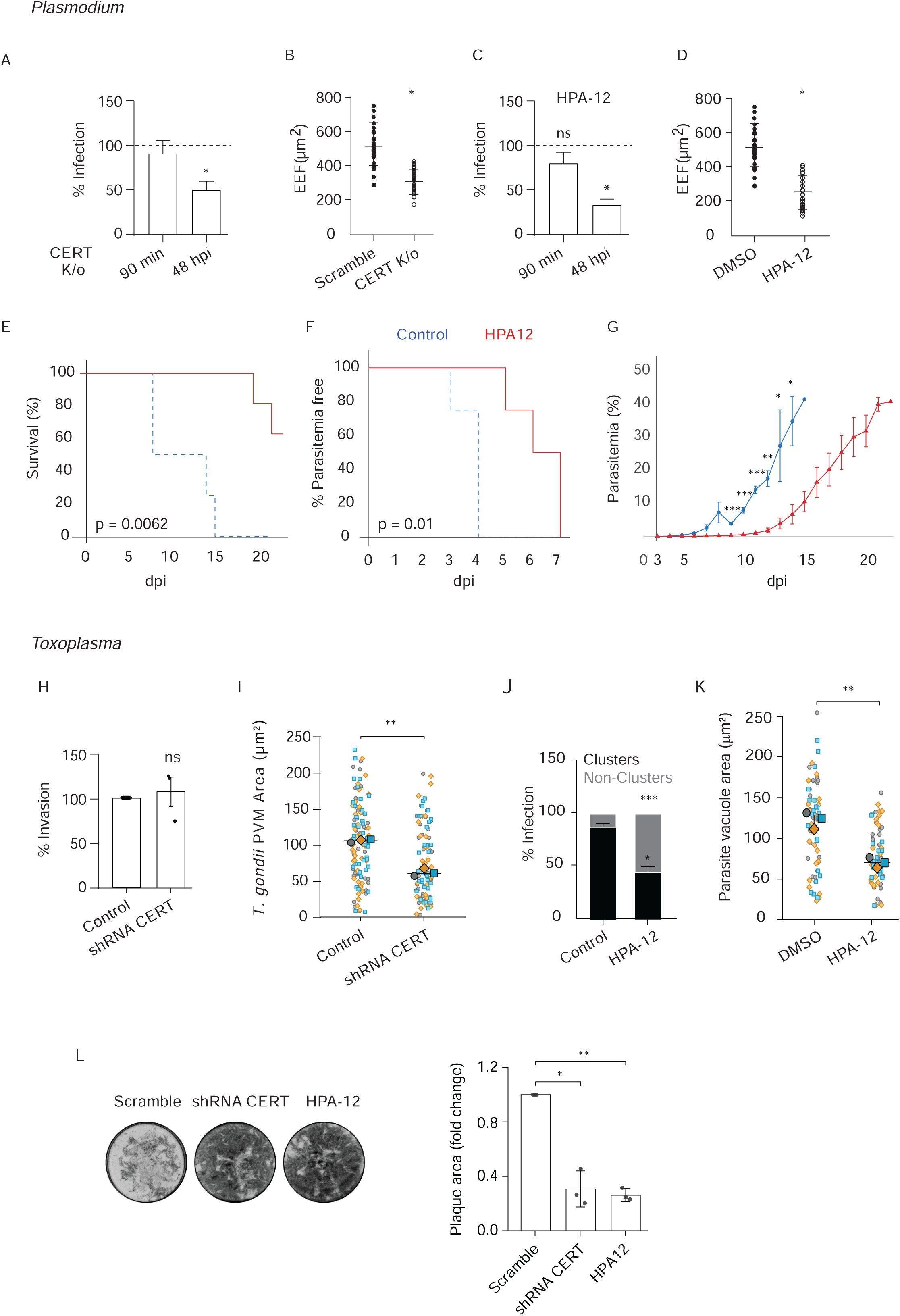
CERT-dependent ceramide transport is required for apicomplexan intracellular development and in vivo growth. **(A-D)** *P. yoelii* liver-stage development in control and CERT knockout (CERT K/o) host cells or cells treated with HPA-12 or vehicle. **(A)** Infection rate at 90 min (invasion) and 48 hpi (development) in CERT K/o host cells. **(B)** Exo-erythrocytic form (EEF) size at 48 hpi in CERT K/o host cells. **(C)** Infection rate at 90 min (invasion) and 48 hpi (development) in HPA-12-treated host cells. **(D)** EEF size at 48 hpi in HPA-12-treated host cells. **(E-G**) *In vivo* infection dynamics in mice infected with *P. berghei* sporozoites and treated with HPA-12 or vehicle. **(E)** Kaplan-Meier survival analysis**, (F)** time to onset of detectable parasitemia, and **(G)** parasitemia progression. Survival analysis was performed using the log-rank (Mantel-Cox) test, with *p* = 0.0062. Time to onset of parasitemia was analyzed using a log-rank (Mantel-Cox) test, with *p* = 0.01. **(H-L)** *T. gondii* fitness following host CERT depletion or HPA-12 treatment. **(H)** Parasite invasion efficiency, **(I)** *T. gondii* normalised mean fluorescence intensity, **(J)** parasite cluster distribution (30 hpi), **(K)** total vacuole area (40 hpi), and **(L)** representative plaque images with plaque area quantification. Data represent mean ± SEM from *n* = 3 independent experiments. * *p* < 0.05, ** *p* < 0.01, *** *p* < 0.001; *P* > 0.05; ns, not significant. Statistical significance for quantitative comparisons was determined using either two-tailed one-sample *t*-test **(H and L)** or Welch’s *t*-test.

**Figure 3.**
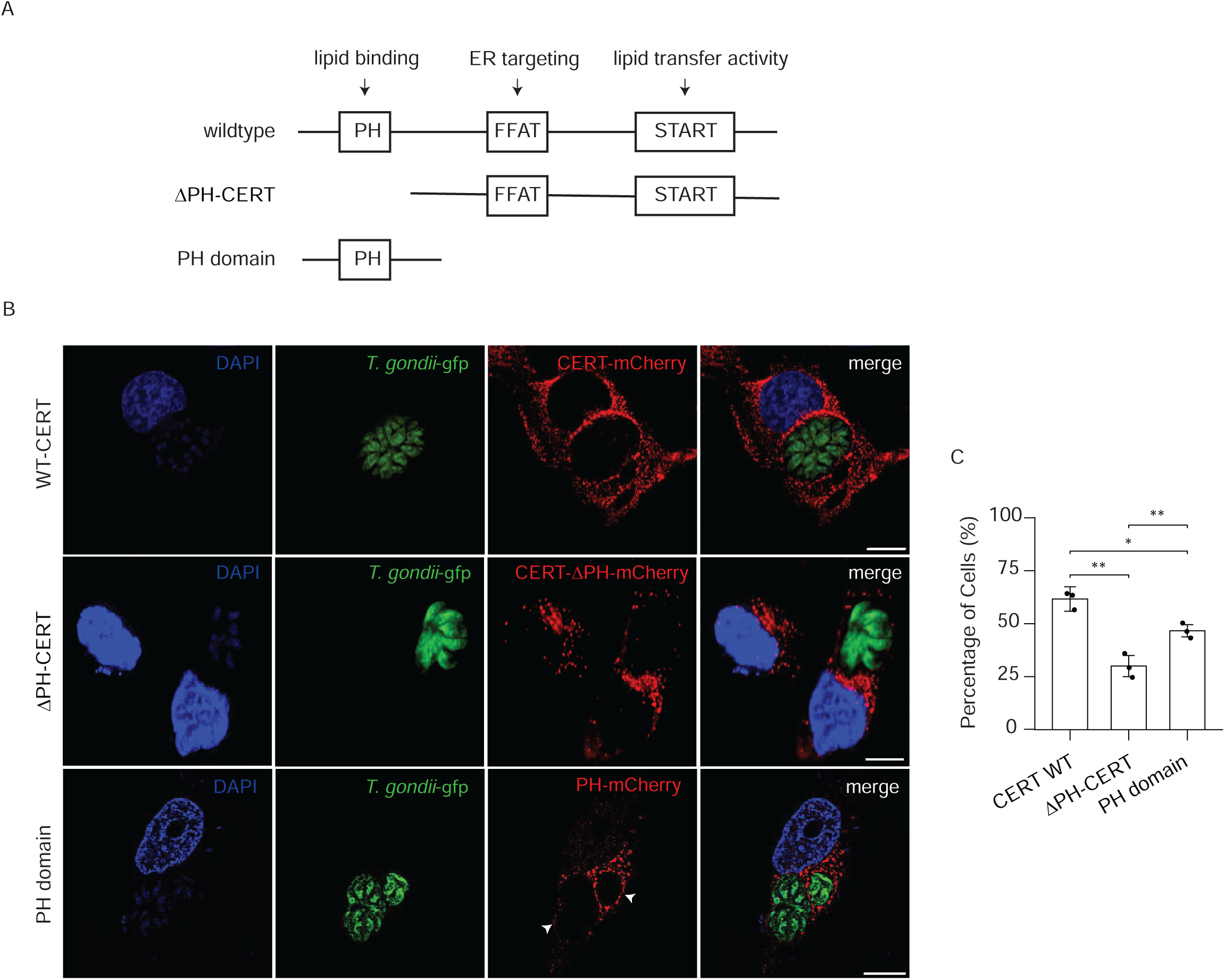
The PH domain of CERT is necessary and sufficient for vacuolar targeting. **(A)** Schematic of CERT domain architecture showing the N-terminal pleckstrin homology (PH) domain, central FFAT motif, and C-terminal START domain, together with the mCherry-tagged constructs used in this study: wild-type CERT (WT), PH domain deletion mutant (ΔPH), and isolated PH domain. **(B)** Representative confocal images of GFP-expressing *T. gondii*-infected host cells expressing WT CERT-mCherry, ΔPH CERT-mCherry, or the isolated PH-mCherry domain. **(C)** Quantification of the percentage of infected host cells showing PVM enrichment for each construct. More than 60 vacuoles were quantified per condition across n = 3 independent experiments. Data represent mean ± SD from *n* = 3 independent experiments. ** *p* < 0.001. Statistical significance was determined using a two-tailed student’s *t*-test. Scale bars, 5 μm.

We next asked whether host CERT-dependent ceramide transport contributes to parasite expansion *in vivo*. Mice treated with HPA-12 displayed a significant delay in the onset of parasitemia, remaining parasitemia-free for an extended period relative to control-treated animals (**Fig. 2F**). Following patency, HPA-12 treatment attenuated the progression of parasitemia (**Fig. 2G**), resulting in significantly prolonged survival of infected hosts (**Fig. 2E**). Thus, pharmacological inhibition of host CERT-dependent ceramide transport restricts *Plasmodium* expansion *in vivo* and delays disease progression.

We next asked whether this functional dependency extends to *T. gondii*. We evaluated parasite fitness following host CERT knockdown (CERT K/d) and HPA-12 treatment. Knockdown of host CERT did not alter *T. gondii* invasion efficiency (**Fig. 2H**), but significantly reduced intracellular parasite size (**Fig. 2I**). Similarly, HPA-12 treatment shifted the distribution of infected cells from replicating parasite clusters toward non-replicating clusters (≤ 2 parasites) (**Fig. 2J**) and significantly decreased total parasite vacuole area (**Fig. 2K**). Furthermore, long-term lytic cycle assays demonstrated a marked reduction in plaque formation under both CERT knockdown and HPA-12 conditions compared with scramble controls (**Fig. 2L**).

Together, these results demonstrate that host CERT-dependent ceramide transport is dispensable for initial parasite invasion but supports intracellular vacuole growth and developmental propagation in both *Plasmodium* and *Toxoplasma*. The *Toxoplasma* and *in vivo Plasmodium* results extend the role of host CERT-dependent lipid transport beyond the previously established *Plasmodium* system and identify CERT-dependent ceramide transport as a shared host requirement for apicomplexan intracellular development.

### The pleckstrin homology domain is necessary and sufficient for CERT recruitment to the parasitophorous vacuole

Having established that host CERT is required for intracellular parasite development, we next sought to define the molecular targeting determinants that recruit CERT to the parasitophorous vacuole. CERT contains three distinct functional modules: an N-terminal pleckstrin homology (PH) domain involved in phosphoinositide lipid binding, a central FFAT motif that mediates ER association through VAP proteins, and a C-terminal START domain responsible for ceramide transfer (**Fig. 3A**). To determine whether vacuolar recruitment depends on the PH domain, we generated mCherry-tagged constructs encoding full-length wild-type CERT (WT), a PH domain deletion mutant (ΔPH-CERT), and the isolated PH domain (**Fig. 3A**).

Confocal imaging of *T. gondii*-infected host cells revealed distinct spatial distributions among the constructs (**Fig. 3B**). Wild-type CERT-mCherry formed a prominent ring around GFP-expressing parasite vacuoles. In contrast, deletion of the PH domain markedly reduced vacuolar enrichment, with ΔPH-CERT showing predominantly diffuse intracellular localization (**Fig. 3B**). In contrast, the isolated PH domain retained clear enrichment at the parasitophorous vacuole interface (**Fig. 3B**).

Quantitative analysis confirmed that wild-type CERT localized to the PVM in the infected cells, whereas deletion of the PH domain reduced PVM localization (**Fig. 3C**). The isolated PH domain was sufficient to confer vacuolar localization (**Fig. 3C**). Thus, although the isolated PH domain does not reproduce the full efficiency of wild-type CERT recruitment, it retains substantial PVM-targeting activity.

Together, these results establish that the PH domain is necessary and sufficient for efficient recruitment of CERT to the parasite vacuole. These findings further distinguish the spatial targeting function of the PH domain from the FFAT-dependent ER association and START-dependent ceramide transfer activities of CERT, identifying the PH domain as the principal determinant that anchors CERT at the host-parasite interface.

### Host VAP proteins are recruited to the parasitophorous vacuole and support apicomplexan development

Having established that CERT is recruited to the parasitophorous vacuole through its PH domain, we next investigated whether ER-resident VAP proteins, canonical organizers of membrane contact sites that interact with FFAT-containing factors, are also positioned at the host-parasite interface. Immunofluorescence microscopy revealed pronounced enrichment of endogenous VAP-B around the parasitophorous vacuole in both liver-stage *P. yoelii* and *T. gondii* infections, forming a distinct envelope encompassing the PVM (**Fig. 4A**). To further examine VAP-B recruitment kinetics during *T. gondii* infection, host cells expressing HA-tagged VAP-B were infected and monitored across development. HA-VAP-B accumulated at the vacuolar interface as early as 4 h post-infection (hpi) and remained stably associated through 44 hpi (**Fig. 4B**). Quantitative analysis confirmed that HA-VAP-B was enriched at the PVM in over 80% of infected host cells (**Fig. 4C**).

**Figure 4.**
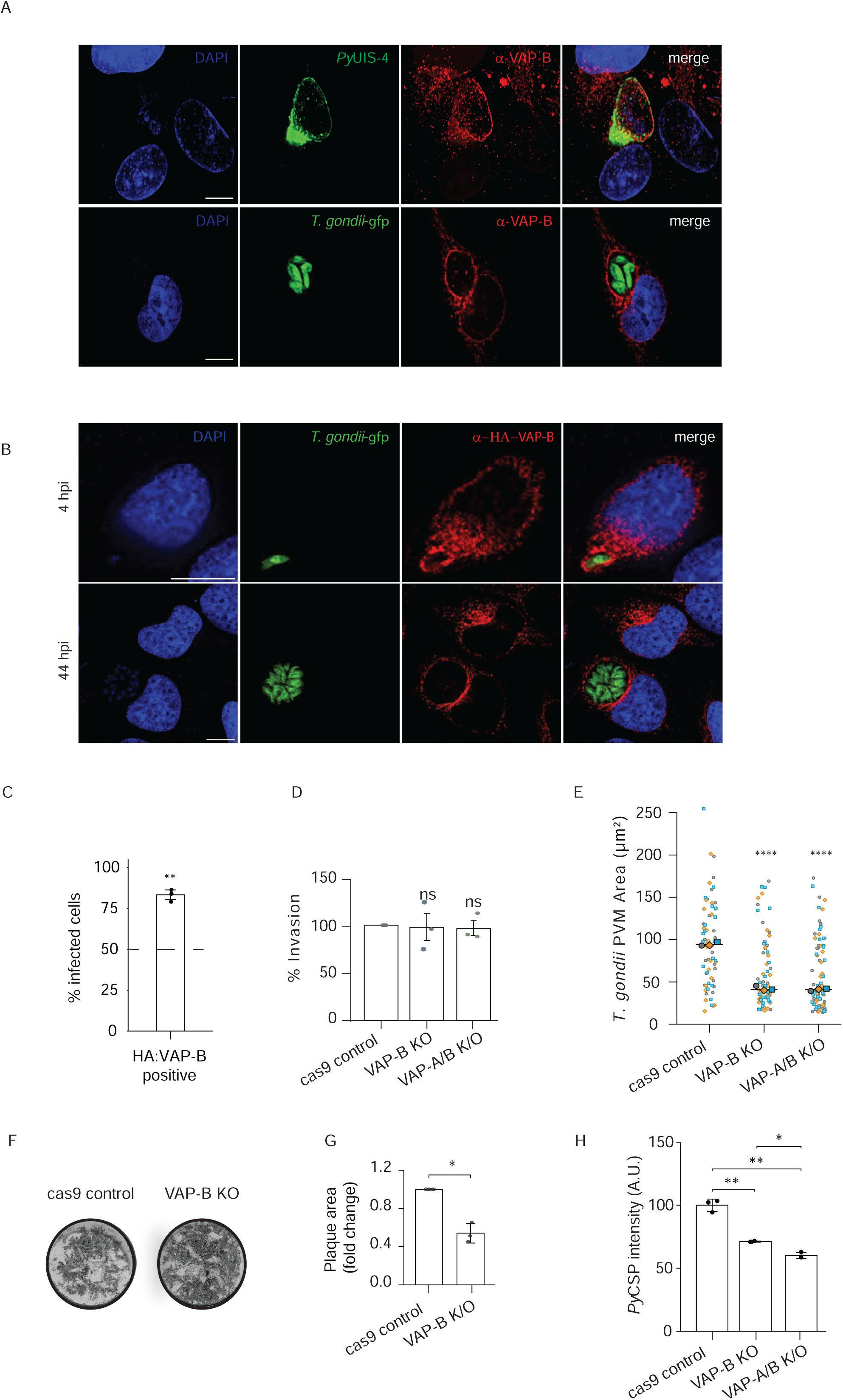
Host VAP proteins are recruited to the vacuole and support apicomplexan development. **(A)** Immunofluorescence images showing endogenous VAP-B enrichment at the parasite vacuole in *P. yoelii*- and *T. gondii*-infected host cells. *P. yoelii* vacuoles are co-stained with PyUIS-4. **(B)** Confocal images showing HA-VAP-B recruitment to the *T. gondii* vacuole at early (4 hpi) and late (44 hpi) intracellular development stages. **(C)** Quantification of the percentage of infected host cells displaying HA-VAP-B enrichment at the *T. gondii* PVM. **(D)** *T. gondii* invasion efficiency in Cas9 control, VAP-B KO, and VAP-A/B double-KO host cells. **(E)** Quantification of *T. gondii* intracellular vacuole area (PVM area, μm²) at 48 hpi in Cas9 control, VAP-B KO, and VAP-A/B double-KO host cells. **(F)** Representative plaque assay images following *T. gondii* lytic cycle growth in Cas9 control and VAP-B KO host cells. **(G)** Quantification of *T. gondii* plaque area fold-change in Cas9 control and VAP-B KO host cells. **(H)** Quantification of *P. yoelii* liver-stage parasite burden based on PyCSP fluorescence intensity in Cas9 control, VAP-B KO, and VAP-A/B double-KO host cells. Data represent mean ± SD or mean ± SEM (**E**) from *n* = 3 independent experiments. *p* < 0.05, *p* < 0.01, *p* < 0.001; ns, not significant (*p* ≥ 0.05). Statistical significance for quantitative comparisons was determined using either two-tailed one-sample *t*-test **(C, D and G)** or Welch’s *t*-test. Scale bars, 5 μm.

To determine whether host VAP proteins are functionally required for parasite development, we generated a single VAP-B knockout (KO; Fig. S2B), shRNA-mediated VAP-A knockdown (Kd; Fig. S2C), and double VAP-A/B KO (Fig. S2B) HeLa cells. In *T. gondii*, neither single nor double VAP knockout affected parasite invasion efficiency (**Fig. 4D**). However, loss of host VAP-B or VAP-A significantly impaired intracellular growth, causing a pronounced reduction in *T. gondii* vacuole area that was similarly observed in double VAP-A/B KO host cells (**Fig. 4E**, Fig. S2D, E). Consistent with a developmental defect, long-term lytic cycle assays demonstrated a marked reduction in plaque area when parasites were cultured in VAP-B KO host cells compared with Cas9 controls (**Fig. 4F, G**).

We further evaluated the requirement for host VAP proteins during liver-stage *Plasmodium* infection. Quantification of *P. yoelii* development by *Py*CSP signal intensity revealed significant reductions in parasite burden in both VAP-B KO and VAP-A/B KO host cells compared with control lines (**Fig. 4H**).

Together, these findings establish host VAP-A and VAP-B as functional components of the ER-PVM interface that support intracellular growth across apicomplexan species. Crucially, while these data demonstrate that VAP proteins are recruited to and required for parasite development, they do not establish direct lipid transfer. Instead, they position host VAPs as candidate ER-side docking platforms, setting up the key question of which parasite-encoded factor engages VAP-B to convert this structural interface into a functional lipid-transfer contact site.

### CERT and VAP-B are required for host sphingolipid enrichment at the parasitophorous vacuole

Having established that CERT and VAP-B accumulate at the parasite vacuole and support intracellular parasite development, we next asked whether these host factors contribute to the enrichment of host-derived sphingolipids at the parasitophorous vacuole membrane (PVM). To track sphingolipid recruitment, host cells were pre-loaded with fluorescent sphingosine prior to *T. gondii* infection. This approach allows the probe to enter the ER-resident host sphingolipid biosynthetic pathway before parasite invasion. In control cells (Scramble and Cas9 control), sphingosine-derived fluorescence concentrated prominently around the parasite vacuole, forming a sharp fluorescent rim outlining GFP-expressing parasites (Fig. 5A, arrowheads). In contrast, shRNA-mediated depletion of host CERT (shRNA CERT) or pharmacological inhibition of CERT using HPA-12 nearly abolished this vacuolar enrichment, yielding a diffuse cytosolic signal (**Fig. 5A**).

**Figure 5.**
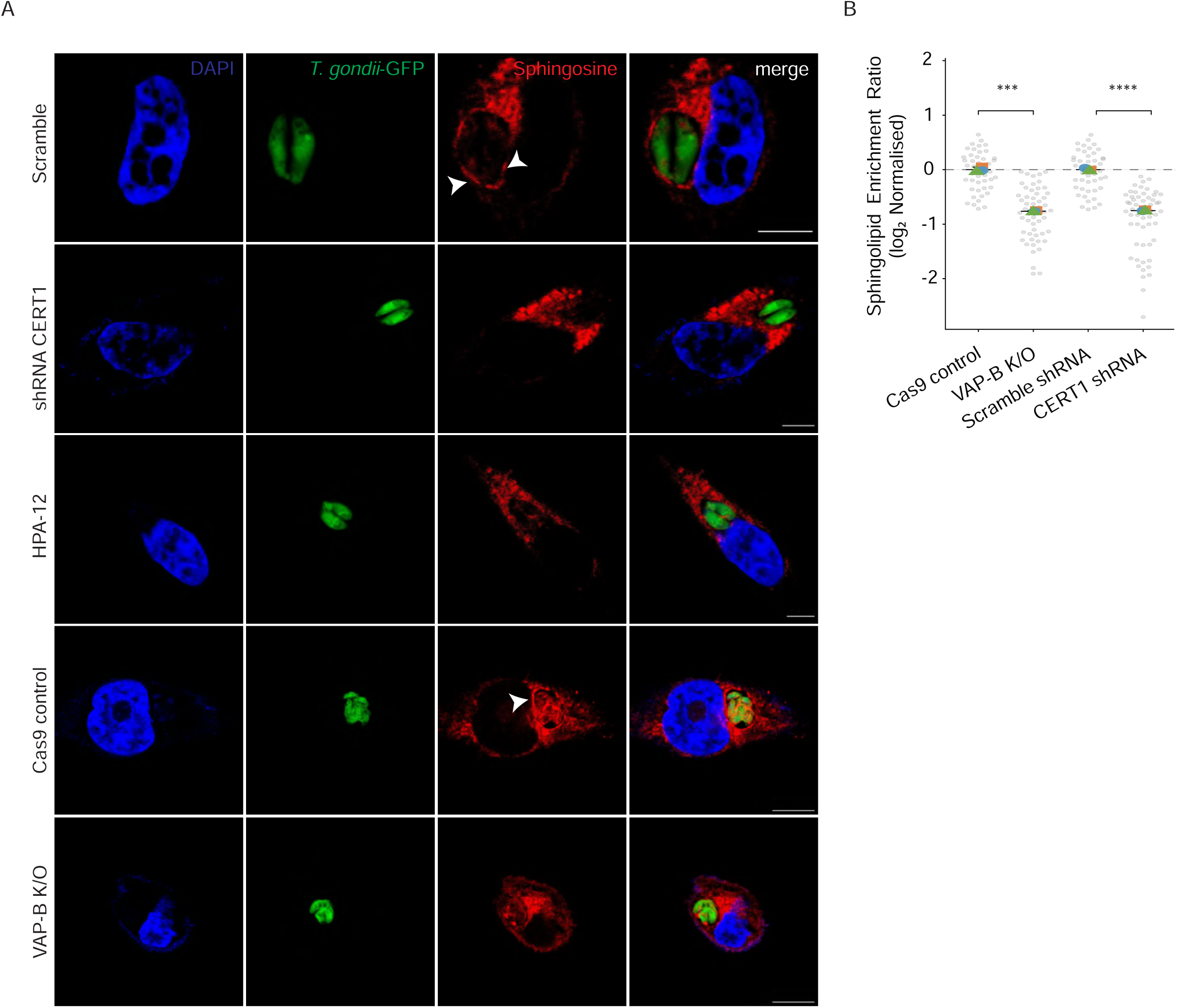
CERT and VAP-B differentially contribute to host sphingolipid enrichment at the parasitophorous vacuole. **(A)** Representative confocal images of host cells pre-labeled with fluorescent sphingosine before *T. gondii* infection under scramble control, Cas9 control, CERT knockdown, HPA-12-treated, and VAP-B knockout conditions. Arrowheads indicate sphingolipid enrichment at the parasite vacuole. **(B)** Quantification of log_2_-normalized vacuolar sphingolipid enrichment ratios relative to the surrounding host cytoplasm. More than 50 vacuoles were analyzed per condition across *n* = 3 independent experiments. Data represent mean ± SD. \**P* < 0.05, ** *p* < 0.01, *** *p* < 0.001; *p* > 0.05; ns, not significant. Statistical significance was determined using a two-tailed Welch’s *t*-test. Scale bar, 5 μm.

We next evaluated whether host VAP-B is similarly required for sphingolipid enrichment at the vacuole. Disruption of host VAP-B (VAP-B K/O) markedly reduced the intense vacuolar enrichment observed in Cas9 control cells (**Fig. 5A, arrowheads**). Quantification of the log-normalized sphingolipid enrichment ratio confirmed statistically significant reductions in vacuolar lipid accumulation following both CERT knockdown and VAP-B knockout (**Fig. 5B**).

Notably, the extent of sphingolipid depletion revealed a clear distinction between CERT and VAP-B perturbations. CERT depletion or inhibition essentially eliminated sphingosine-derived enrichment at the parasite vacuole, whereas VAP-B knockout produced a substantial but incomplete reduction (**Fig. 5A, B**). This differential phenotype indicates that sphingolipid enrichment at the PVM is strongly dependent on CERT, whereas VAP-B contributes to the efficiency of this process.

Together, these findings demonstrate that CERT and VAP-B contribute differentially to host sphingolipid enrichment at the *T. gondii* vacuole. The incomplete loss of sphingolipid enrichment following VAP-B depletion is consistent with a model in which CERT can access host sphingolipid pools independently of VAP-B, while VAP-B enhances the efficiency of CERT-dependent lipid delivery. These functional results prompted us to identify the parasite-encoded machinery that recruits VAP-B and establishes the host-parasite contact interface.

### TgVPS13 is a parasite-encoded FFAT effector that physically interacts with host VAP-B at the ER-PVM interface

Having established that CERT and VAP-B contribute to host sphingolipid accumulation at the *T. gondii* vacuole, we next sought to identify the parasite-encoded machinery that engages host VAP-B at the ER-PVM interface. Because VAP proteins recognize FFAT-containing proteins at membrane contact sites, we integrated in situ VAP-B proximity labeling of VAP-B (**Fig. S3A, B, C, D and E; Supplementary Data 2**) with bioinformatic FFAT motif predictions using the scoring framework described by Slee and Levine^17^. Plotting log_2_ fold change (VAP-B/NES) against predicted FFAT scores recovered canonical host VAP interactors, including CERT1 and OSBP1, alongside the recently identified parasite effector TgVIP1, validating the screen (**Fig. 6A).** Intersecting the 144 parasite specific hits (log_2_ fold change ≥ 1) with proteins containing high-confidence FFAT motifs (FFAT motif score < 2) yielded 4 candidate hits (Fig. 6B). Among the top three (excluding RNA dependent RNA polymerase; TGGT_217182) candidate parasite proteins (TGGT1_210700, TGGT1_306020 and TGGT1_212880), TGGT1_210700 emerged as the top candidate based on its predicted transmembrane domain and sequence similarity to the VPS13 family of VAP-binding lipid transfer proteins, leading us to designate it TgVPS13 (**Fig. 6A, B**).

**Figure 6.**
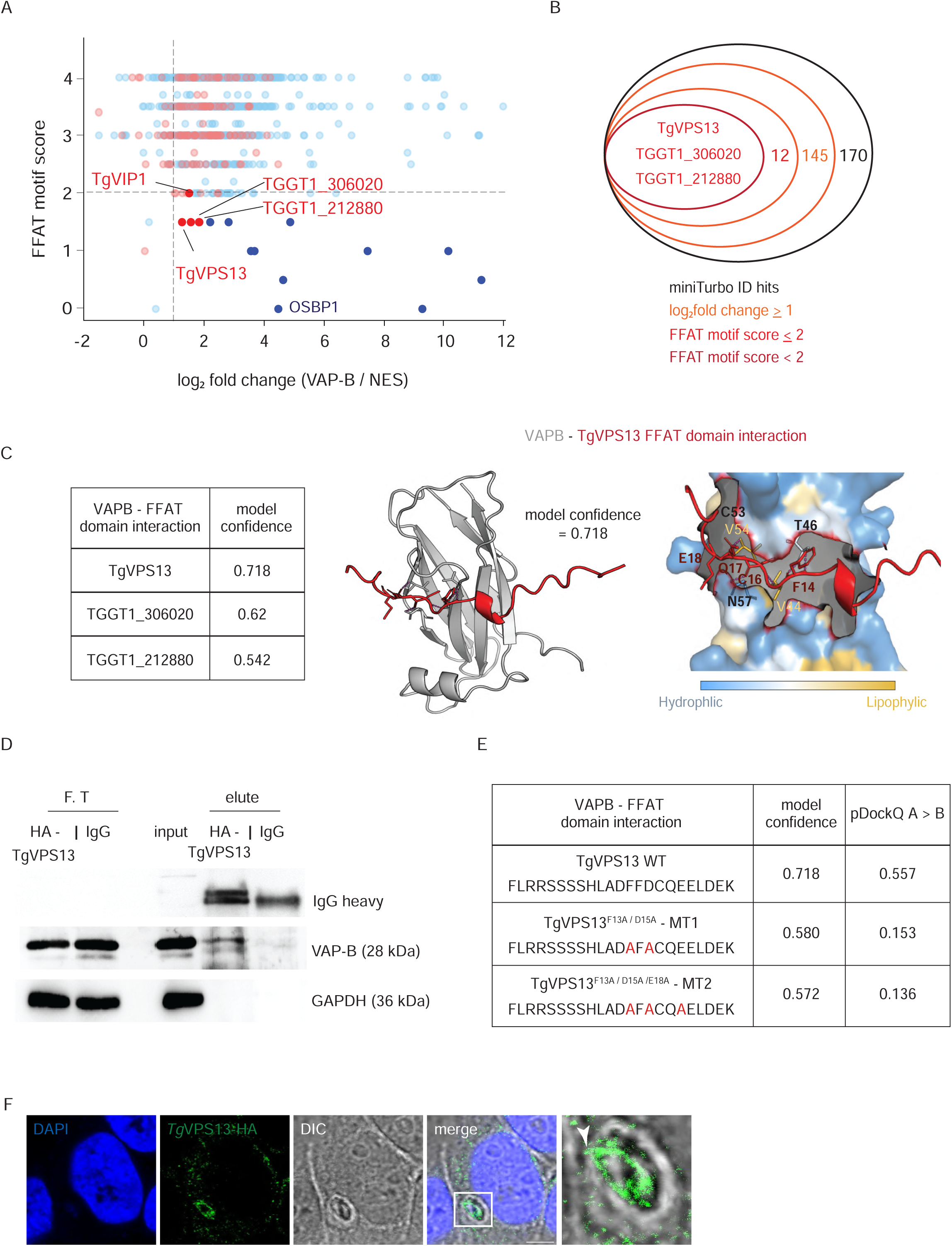
TgVPS13 is a parasite-encoded FFAT effector that physically interacts with host VAP-B. **(A)** Integration of host VAP-B miniTurbo proximity labeling with bioinformatic FFAT motif prediction scores. Canonical host VAP interactors (CERT1, OSBP1) and parasite effector TgVIP1 validate the screen alongside prioritized parasite candidates. **(B)** Nested Venn diagram intersecting miniTurbo proximity hits with predicted high-confidence FFAT motifs, yielding three prioritized parasite candidate hits. **(C)** Structural model of the TgVPS13 FFAT domain bound to the host VAP-B MSP domain (model confidence = 0.718). Ribbon diagram (left) and lipophilicity surface map (right) highlight key contact residues between TgVPS13 (F14, C16, D17, E18) and host VAP-B (T46, C53, N57). **(D)** Co-immunoprecipitation of HA-tagged TgVPS13 from *T. gondii*-infected host cells followed by immunoblotting for endogenous host VAP-B and host GAPDH (loading control). **(E)** *In silico* interaction predictions with host VAP-B. Sequence alignment, AlphaFold-Multimer model confidence, and interface quality metrics (pDockQ A>B) for Wild-Type (WT) TgVPS13 and alanine mutants (MT1: F13A/D15A; MT2: F13A/D15A/E18A), highlighting the collapse in interface quality upon motif mutation. **(F)** Representative confocal immunofluorescence images showing localization of HA-TgVPS13 (green) along the parasitophorous vacuole periphery. Nuclei are counterstained with DAPI (blue); BF, brightfield. Inset provides a zoomed view of vacuolar enrichment (arrowhead). Scale bars, 5 μm.

Structural modeling of the TgVPS13 FFAT sequence with the host VAP-B MSP domain predicted a high-confidence complex (model confidence = 0.718, pDockQ A>B = 0.557; **Fig. 6C, E**). The model positions key hydrophobic and acidic residues of the TgVPS13 FFAT motif (F14, C16, D17, E18) within the canonical binding pocket of the VAP-B MSP domain, forming specific contacts with host residues V44, T46, V54, C53, and N57 (**Fig. 6C; Supplementary data 3**). To test the structural dependency of this interaction on conserved motif residues, we performed *in silico* alanine mutagenesis on key positions within the FFAT core (**Fig. 6C**). Alanine substitutions in Mutant 1 (F13A/D15A) and Mutant 2 (F13A/D15A/E18A) led to a drop in overall model confidence (0.580 and 0.572, respectively) and a severe collapse in interface quality, with pDockQ A>B scores dropping sharply from 0.557 in WT to 0.153 in Mutant 1 and 0.136 in Mutant 2 (**Fig. 6C; Supplementary data 3**). This precipitous decrease in pDockQ A>B demonstrates that these core residues are essential for maintaining the binding interface with host VAP-B.

To verify this physical association in infected host cells, we performed immunoprecipitation assays using HA-tagged TgVPS13. Immunoprecipitation of HA-TgVPS13 specifically co-purified endogenous host VAP-B (28 kDa), whereas no VAP-B signal was detected in the IgG control, and host GAPDH (36 kDa) was cleanly excluded from the eluate (**Fig. 6D**).

In line with a functional role at the host-parasite interface, confocal immunofluorescence microscopy demonstrated that HA-tagged TgVPS13 accumulates along the parasitophorous vacuole periphery (**Fig. 6F**).

### TgVPS13 is required for vacuolar sphingolipid enrichment, host CERT recruitment, and parasite intracellular development

To evaluate the spatial organization of TgVPS13 in relation to host contact site machinery, we performed confocal immunofluorescence microscopy. Co-staining demonstrated distinct co-localization between host VAP-B and HA-tagged TgVPS13 along the PVM (**Fig. 7A**).

**Figure 7.**
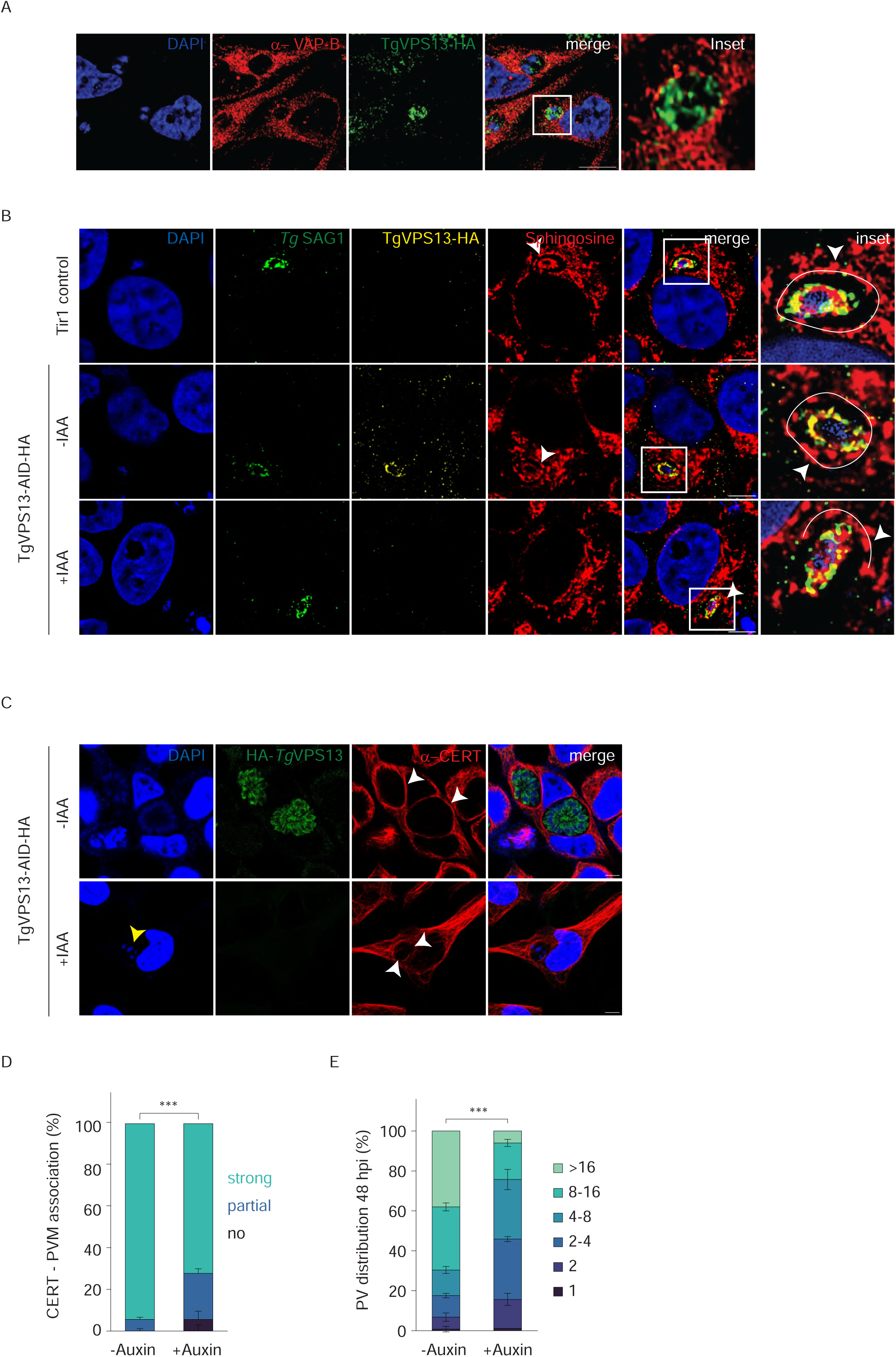
TgVPS13 is required for vacuolar sphingolipid enrichment, host CERT recruitment, and parasite intracellular development. **(A)** Representative confocal immunofluorescence microscopy showing spatial co-localization of host VAP-B (red) and HA-TgVPS13 (green) at the parasitophorous vacuole membrane (PVM). Inset highlights spatial association at the host-parasite interface. Nuclei are counterstained with DAPI (blue). **(B)** Representative confocal images of host pre-labeled fluorescent sphingosine (red) in Tir1 control, untreated TgVPS13-AID-HA (-IAA), and auxin-treated TgVPS13-AID-HA (+IAA) parasites. Parasites are identified by TgSAG1 (green) and TgVPS13-AID-HA (yellow). White outlines delineate the PVM boundary (inset, arrowheads), showing selective loss of PVM sphingosine concentration upon TgVPS13 degradation while intraparasitic signal remains intact. **(C)** Confocal immunofluorescence microscopy of host CERT (red) recruitment in TgVPS13-AID-HA parasites ± IAA. TgVPS13 depletion (+IAA) causes partial loss of host CERT accumulation along the PVM (white arrowheads) and results in disoriented intravacuolar parasite organization and disarranged nuclear architecture (yellow arrowhead). (D) Quantification of CERT-PVM association percentage categories (strong, partial, no association) across treatment groups. Error bars represent single-sided standard error of the mean (SEM) across three distinct biological replicates. Statistical divergence between overall phenotypic class distributions was evaluated via a Chi-square test of independence (*** *p* < 0.001). **(E)** Quantification of parasitophorous vacuole (PV) size distribution (parasites per PV) at 48 hours post-infection (hpi) in TgVPS13-AID-HA parasites ± IAA (*** *p* < 0.001). Statistical significance between overall replication profiles was determined by two-sided Pearson’s Chi-square test on pooled vacuole counts (χ² = 121.2, df = 5, *** *p* < 0.001, total *N* = 518 PVs). Error bars represent single-sided standard error of the mean (SEM) across three distinct biological replicates.

To test whether TgVPS13 is functionally required for sphingolipid enrichment at the vacuole, we generated an auxin-inducible degron parasite strain (TgVPS13-AID-HA). In Tir1 control cells and untreated TgVPS13-AID-HA (-IAA) parasites, fluorescent sphingosine was markedly enriched along the parasitophorous vacuole membrane (delineated by the white PVM boundary outline in **Fig. 7B**, arrowheads). Upon auxin-induced degradation (TgVPS13-AID-HA; +IAA), TgVPS13-HA was efficiently depleted, resulting in a pronounced loss of sphingosine accumulation within the outlined PVM boundary (**Fig. 7B**). Notably, internal intra-parasitic sphingosine signal as well as host cellular fluorescence remained completely unchanged, demonstrating that TgVPS13 specifically drives localized sphingolipid accumulation at the ER-PVM interface rather than overall parasite lipid uptake (**Fig. 7B)**.

Because TgVPS13 engages host contact site machinery, we next evaluated whether its depletion impacts host CERT organization at the ER-PVM interface. In untreated TgVPS13-AID-HA (-IAA) parasites, TgVPS13-HA localized along the vacuolar periphery in close association with recruited host CERT (**Fig. 7C**, white arrowheads). Upon TgVPS13 depletion (+IAA), CERT accumulation at the PVM was partially reduced (**Fig. 7C**, white arrowheads). Quantitative evaluation of CERT-PVM association confirmed significant reduction in strong CERT recruitment upon TgVPS13 degradation (**Fig. 7D**). The partial association observed in a subset of TgVPS13-depleted vacuoles is consistent with the intrinsic ability of CERT’s PH domain to recognize PVM-associated lipids, whereas the loss of the TgVPS13-VAP-B interaction prevents maximal CERT concentration at host-parasite contact sites.

In addition to impaired lipid machinery recruitment, loss of TgVPS13 led to visible defects in parasite structural integrity. Depleted vacuoles (+IAA) exhibited disoriented parasite organization and disarranged nuclear architecture, as highlighted by aberrant DAPI structures (**Fig. 7C**, yellow arrowhead). Consistent with these morphological and metabolic defects, quantification of parasitophorous vacuole (PV) size distribution at 48 hours post-infection (hpi) demonstrated a severe replication defect upon TgVPS13 depletion, evidenced by a significant accumulation of smaller vacuoles containing 1-4 parasites per PV compared to untreated (-IAA) controls (*p* < 0.001) (**Fig. 7E**).

Together, these findings establish TgVPS13 as a parasite-encoded FFAT effector that engages host VAP-B and promotes efficient host CERT recruitment and sphingolipid enrichment at the parasitophorous vacuole, driving parasite intracellular replication.

### Discussion

Membrane contact sites (MCSs) are increasingly recognized as essential platforms for inter-organelle communication^14^, yet their functional significance during intracellular infection has remained poorly defined^13,18^. In *T. gondii*, proximity-labeling approaches first identified host and parasite proteins at the parasitophorous vacuole membrane (PVM), including MOSPD2 and other components of the host-parasite interface^13,19^. Subsequent studies established host VAP proteins and MOSPD2 as components of ER-PV interactions ^11–13^ and identified parasite FFAT-like proteins, including *Tg*VIP1^12^ and *Tg*ROP1^11^, that engage host VAP proteins to establish physical contact. In parallel, host CERT has been implicated in *Plasmodium* liver-stage development and ceramide acquisition^7,8^. Whether ER-PVM contacts functionally connect to host lipid-transfer machinery, however, remained unresolved. Here, we establish this connection by showing that host CERT mediates sphingolipid enrichment at the PVM, while the parasite-encoded FFAT effector TgVPS13 engages host VAP-B to spatially organize this lipid-transfer pathway.

Host-derived lipids are critical resources for intracellular apicomplexans^2–5,8,20,21^, but how specific host lipid pools are mobilized to the PVM has remained a central question. Our finding that fluorescent sphingosine pre-loaded into host cells subsequently enriches around intracellular *T. gondii* indicates that parasites access pre-existing host sphingolipid pools that have entered the endogenous biosynthetic pathway. This observation complements recent work showing that sphingolipids support *Plasmodium* liver-stage development^2^ and that host CERT contributes to ceramide trafficking to the parasite vacuole^8^. Because ceramide biosynthesis is concentrated at the host ER^6,22^, the close physical association between host ER networks and the PVM^13,18^ provides a spatial solution to the challenge of accessing this lipid pool. Recent ultrastructural analysis further demonstrates that the PVM itself has an unusual lipid organization^23^. Together, these observations support the view that the PVM is not simply a passive membrane surrounding the parasite, but a specialized lipid interface whose composition and organization are actively remodeled during infection. Our findings extend this concept by identifying a mechanism through which host-derived sphingolipids are delivered to this specialized membrane.

Host CERT emerges as the central component of this pathway^6,9,10^. Genetic depletion and pharmacological inhibition of CERT severely impaired intracellular development in both P. *yoelii* and *T. gondii* without perturbing invasion, indicating that CERT-dependent lipid transport is required during intracellular parasite proliferation. Importantly, domain-mapping experiments revealed that the N-terminal PH domain of CERT is necessary and sufficient for vacuolar targeting^10,24^, providing a mechanism for positioning the lipid-transfer machinery directly at the membrane onto which ceramide is deposited. These findings extend the previously described role of CERT in *Plasmodium^7,8^* by demonstrating that PVM targeting of CERT and its requirement for intracellular development are conserved in *T. gondii*.

The differential effects of CERT and VAP-B perturbation provide a novel mechanistic insight into how this contact site functions. CERT depletion or inhibition nearly abolished sphingolipid enrichment at the PVM, whereas VAP-B loss produced a substantial but incomplete reduction. Thus, CERT represents the principal and indispensable component for sphingolipid accumulation, whereas VAP-B enhances the efficiency of this process. Importantly, VAP-B loss does not phenocopy CERT loss, indicating that CERT can access the PVM independently of VAP-B. Instead, VAP-B-dependent ER-PVM proximity appears to increase the efficiency with which PVM-associated CERT accesses its ER-derived lipid pool. This distinction is important in light of recent studies showing that VAP-dependent contacts are major determinants of ER-PV architecture^11–13^. Rather than functioning as an obligatory transport conduit, the MCS may therefore act as a spatial amplifier that concentrates the appropriate membranes and transfer machinery to enhance lipid flux.

TgVPS13 provides a parasite-side mechanism for establishing this spatial organization. TgVPS13 emerged from the VAP-B proximity proteome as a parasite protein containing a high-confidence FFAT motif and was independently supported by co-immunoprecipitation and vacuolar localization as a VAP-B-associated factor^25,26^ including recently discovered, TgVIP1^12^. Structural modeling of its FFAT sequence with the VAP-B MSP domain further supported the predicted interaction, while mutation of conserved FFAT^11,27,28^ residues markedly reduced the predicted interface quality. These structural and biochemical observations are complemented by functional experiments showing that conditional depletion of TgVPS13 reduces PVM sphingolipid enrichment, impairs CERT recruitment and compromises intracellular parasite development. A particularly informative feature of this phenotype is its spatial specificity: TgVPS13 depletion markedly reduced sphingosine enrichment along the PVM, whereas the internal sphingosine signal within the parasite remained unchanged. The residual CERT association with the PVM following TgVPS13 depletion is consistent with the intrinsic ability of the CERT PH domain to recognize PVM-associated lipids independently of VAP-B. Together, these findings identify TgVPS13 as a parasite factor that links VAP-dependent ER-PVM organization to localized host sphingolipid acquisition, enhancing rather than absolutely determining CERT-dependent lipid transfer at the host-parasite interface (**Fig. 8**).

**Figure 8:**
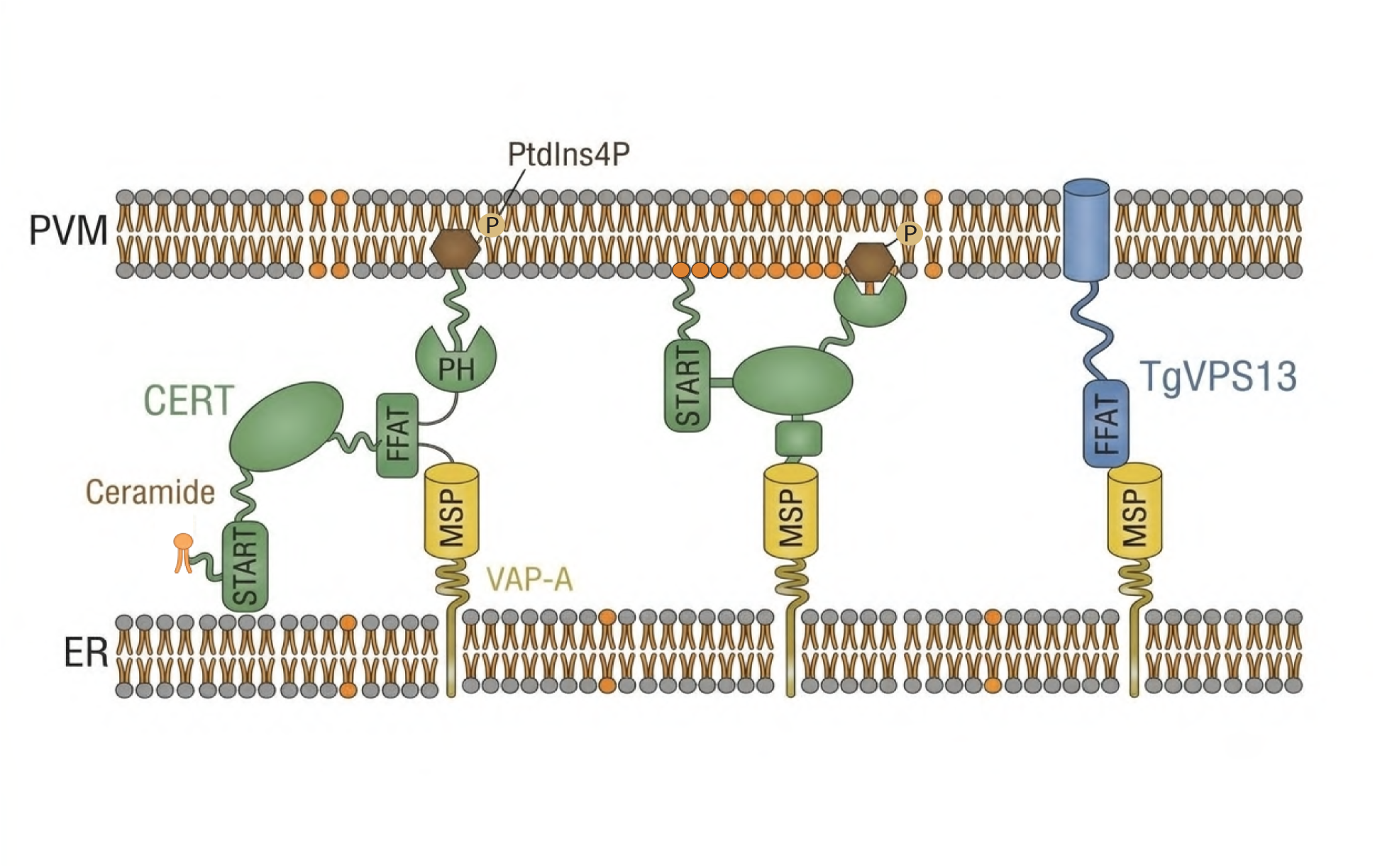
Model for *Tg*VPS13-VAP-B-mediated ER-PVM proximity and CERT-dependent ceramide delivery. Schematic model illustrating the proposed mechanism of host ceramide acquisition at the ER-PVM interface. Host CERT is recruited to the PVM through its PH domain, positioning its START domain at the vacuolar membrane. Parasite-encoded *Tg*VPS13 engages host VAP-B through its FFAT-containing region, bringing the host ER into proximity with the PVM. This spatial organization enhances access of PVM-associated CERT to ER-derived ceramide and facilitates ceramide deposition onto the PVM, supporting intracellular parasite growth.

More broadly, our findings extend the emerging view of pathogen-induced MCSs from structural interfaces to metabolically functional platforms^11–13,18^. The recent demonstration that the PVM of both *T. gondii* and *P. falciparum* has an unusual lipid organization further emphasizes that this membrane represents a specialized host-parasite interface rather than a conventional host-derived membrane^23^. Our work adds a metabolic dimension to this framework by connecting ER-PVM organization to host sphingolipid acquisition. Whether analogous FFAT-containing parasite factors organize CERT-dependent lipid acquisition during *Plasmodium* infection remains an important question. The conservation of CERT dependence and VAP-associated phenotypes across *T. gondii* and liver-stage *Plasmodium* nevertheless suggests that exploitation of host ER lipid-transfer machinery may represent a broader feature of apicomplexan biology.

Together, our findings establish the ER-PVM contact site as a metabolically functional interface that enhances host sphingolipid acquisition. Rather than evolving an independent lipid-transfer apparatus, *T. gondii* combines PVM-targeted host CERT with parasite-mediated remodeling of ER-PVM proximity through TgVPS13-VAP-B. This provides a spatial solution to the challenge of accessing ER-derived ceramide and illustrates how apicomplexan parasites repurpose host organelle contact-site machinery to acquire essential host metabolites.

### Methods

#### Cell culture and parasite maintenance

Human cervical adenocarcinoma cells (HeLa, ATCC CCL-2) and human foreskin fibroblasts (HFF, ATCC SCRC-1041) were maintained in Dulbecco’s modified Eagle medium (DMEM; Gibco) supplemented with 10% South American fetal bovine serum (SA-FBS; Gibco). HepG2 cells stably expressing CD81 (HepG2-CD81) were cultured in DMEM supplemented with 10% North American fetal bovine serum (NA-FBS; Gibco). All host cell lines were maintained at 37°C in a humidified incubator under 5% CO₂. *Toxoplasma gondii* tachyzoites were propagated by serial passage in HeLa or HFF monolayers. Freshly egressed parasites were isolated by passage through a 27G needle and filtration through a 5-μm polycarbonate membrane before infection assays. Hepatic-stage *Plasmodium* cultures were maintained in HepG2-CD81 cells.

#### *Plasmodium* transmission and sporozoite preparation

All animal procedures were approved by the Institutional Animal Ethics Committee. For generation of infectious sporozoites, 6- to 8-week-old donor mice were inoculated with blood-stage *Plasmodium berghei* ANKA or *Plasmodium yoelii* 17XNL parasites. Following confirmation of functional gametocytes by *in vitro* exflagellation, anesthetized mice were exposed to 3- to 6-day-old female *Anopheles stephensi* mosquitoes. Blood-fed mosquitoes were maintained at 75% relative humidity under a 12-h light/dark cycle and supplied with 10% sucrose. *P. berghei*-infected mosquitoes were maintained at 19-20°C, whereas *P. yoelii*-infected mosquitoes were maintained at 21-24°C. Midgut oocyst development was evaluated microscopically at days 7-9 post-infection for *P. yoelii* and days 11-13 for *P. berghei*. Salivary gland sporozoites were harvested at day 14 post-infection for *P. yoelii* and day 20 for *P. berghei*. Surface-sterilized mosquitoes were dissected in ice-cold DMEM, and pooled salivary glands were mechanically disrupted. Sporozoites were quantified using a hemocytometer and used immediately for infection experiments.

#### Genetic manipulation and molecular cloning

All the primer, shRNA and sgRNA guide sequences used in the study are provided in the **Supplementary Data 1**.

#### Generation of CERT knockout HepG2-CD81 cells

GFP-tagged lentiviral vectors carrying gene-specific CRISPR sgRNAs or a scramble control were obtained from ABM Good (Richmond, BC, Canada). Non-replicating lentiviral stocks were generated in HEK293-FT cells using polyethyleneimine MAX (PEI MAX, Polysciences) and third-generation lentiviral packaging mix. Viral supernatants were collected 36 h after transfection and filtered through 0.45-μm syringe filters.

HepG2-CD81 cells were transduced with lentiviral supernatants in the presence of 0.5 μg/mL polybrene (Sigma-Aldrich). Twenty-four hours after transduction, cells were selected with 2 μg/mL puromycin for at least 5 days. For experiments using knockout populations, analyses were restricted to GFP-positive cells. Knockout efficiency was validated by western blot analysis.

#### Generation of VAP-B knockout and shRNA knockdown cell lines

VAP-B knockout was generated in HeLa cells using CRISPR/Cas9-mediated genome editing with the lentiCRISPRv2 vector (Addgene #52961). The vector was linearized with Esp3I, and complementary oligonucleotides encoding VAPB, VAPA/B, or scramble sgRNAs were annealed, phosphorylated, and ligated into the vector using T4 DNA ligase. Positive clones were verified by Sanger sequencing.

For shRNA-mediated knockdown, lentiviral constructs from The RNAi Consortium (TRC) shRNA library were obtained through the shRNA Resource Center at the Indian Institute of Science. Constructs targeting *VAPA* and *CERT1*, together with a non-targeting scramble control, were used. Lentiviral particles were generated in HEK293T cells by transient co-transfection with psPAX2 and pMD2.G packaging plasmids. HeLa cells were transduced in the presence of polybrene and selected with puromycin to establish stable cell populations. Knockout and knockdown efficiencies were validated by western blot analysis.

#### CERT1 overexpression plasmid and domain variant cloning

Full-length human CERT1 and its domain variants were cloned into the mCherry-N1 mammalian expression vector (Addgene #54680). The vector backbone was linearized using EcoRI and KpnI (New England Biolabs), and coding sequences for full-length CERT1, the PH domain, and the ΔPH variant were amplified from HeLa cDNA using construct-specific primers containing terminal restriction sites. PCR products were digested with EcoRI and KpnI and directionally ligated into the mCherry-N1 backbone using T4 DNA ligase. Ligated plasmids were transformed into competent *E. coli*, and positive clones were isolated and verified. Primer sequences are provided in the supplementary primer table.

#### *T. gondii* genome editing and endogenous tagging

For endogenous C-terminal tagging of *T. gondii* VPS13, CRISPR/Cas9-mediated homology-directed repair was used to introduce an auxin-inducible degron (AID) and 3×HA epitope tag at the endogenous locus. A guide RNA targeting the C-terminal region of *TgVPS13* was cloned into pU6-Universal (Addgene #52694) following BsaI-HFv2 digestion and ligation of complementary oligonucleotides. The HDR donor contained the AID-3×HA cassette together with a dihydrofolate reductase (DHFR) selection marker and homology sequences flanking the endogenous stop codon and was amplified from the modified pLinker-AID-3×HA-HXGPRT-LoxP backbone (Addgene #86669).

The sgRNA/Cas9 plasmid and linearized HDR template were co-electroporated into the *T. gondii* RH88 Δ*ku80* strain expressing TIR1. Recombinant parasites were selected in HFF cells using 2.5 μM pyrimethamine for 2 weeks and subsequently used for conditional depletion and localization experiments.

#### Parasite infection and imaging

##### *T. gondii* infection and immunofluorescence microscopy

For transient transfection experiments, host cells were seeded on glass coverslips in 24-well plates and transfected 24 h later using Lipofectamine 3000 (Thermo Fisher Scientific) according to the manufacturer’s instructions. Freshly egressed *T. gondii* tachyzoites were isolated by 27G needle passage and 5-μm filtration, centrifuged at 800 × *g* for 5 min, and added to host monolayers at an MOI of 0.5. Unattached extracellular parasites were removed by washing with PBS at 3 h post-infection.

Cells were fixed with 4% paraformaldehyde (PFA) in PBS for 20 min at room temperature, permeabilized with 0.1% Triton X-100 for 10 min, and blocked with 2% BSA in PBS for 1 h. Primary antibodies were incubated overnight at 4°C, followed by fluorophore-conjugated secondary antibodies for 1 h at room temperature. Nuclei were counterstained with DAPI for 10 min, and coverslips were mounted using Fluoromount-G (SouthernBiotech). Fluorescence datasets were acquired using Olympus BX53, Olympus IX83, Olympus FV3000, or Leica STELLARIS microscopes and processed using Fiji/ImageJ.

##### Flow cytometric analysis of *T. gondii* infection

HeLa cells were infected with eGFP-expressing *T. gondii* tachyzoites for 90 min. Monolayers were harvested by trypsinization, washed with PBS, fixed with 4% PFA for 20 min at room temperature, and analyzed using a BD FACSLyric flow cytometer to determine the percentage of GFP-positive infected cells and mean fluorescence intensity.

##### *Plasmodium* liver-stage infection

HepG2-CD81 wild-type or knockout/knockdown cells were seeded at 5 × 10^5^ cells per well in 24-well plates and infected with *P. yoelii* sporozoites at an MOI of 0.3. For assessment of sporozoite invasion, infections were terminated at 90 min post-infection. For liver-stage development experiments, the medium was replaced after 90 min and infection was allowed to proceed for 48 h. For drug-treatment experiments, cells were either pretreated for 24 h before infection and washed before sporozoite addition or treated after 24 h of infection, as indicated, and infection was allowed to proceed for 48 h.

##### Flow cytometric analysis of *Plasmodium* sporozoite invasion

HepG2-CD81 wild-type or knockout/knockdown cells were infected with *P. yoelii* sporozoites at an MOI of 0.3 for 90 min. Cells were detached using Accutase (Life Technologies), fixed using Cytofix/Cytoperm (BD Biosciences), blocked for 1 h at room temperature in Perm/Wash buffer containing 2% BSA, and stained overnight at 4°C with Alexa Fluor 488- or Alexa Fluor 647-conjugated circumsporozoite protein (CSP) antibody. Cells were washed and resuspended in PBS containing 5 mM EDTA. Infection rates were determined by flow cytometry using an LSR II flow cytometer (BD Biosciences) and analyzed using FlowJo (Tree Star).

##### Immunofluorescence analysis of *Plasmodium* liver-stage infection

For imaging experiments, HepG2-CD81 wild-type or knockout cells were seeded in 8-well chamber slides and infected with *P. yoelii* sporozoites. Cells were fixed with 3.7% (v/v) paraformaldehyde at the indicated time points, including 90 min, 24 h, and 48 h post-infection. Cells were permeabilized with Triton X-100 and stained with fluorescently labeled UIS-4 or other antibodies as indicated. Nuclei were counterstained with DAPI using Vectashield.

##### Image analysis and parasite growth quantification

For *Plasmodium* liver-stage growth measurements, UIS-4 fluorescence was used to delineate the parasitophorous vacuole, and the area enclosed by the UIS-4-positive boundary was quantified using Imaris (Bitplane) to determine the size of individual liver-stage exoerythrocytic forms.

For *T. gondii*, parasite invasion at 90 min post-infection was quantified by flow cytometry using GFP-expressing parasites. For intracellular growth at 48 h post-infection, GFP-positive parasite area was quantified from fluorescence microscopy images using Fiji/ImageJ. For vacuole-area measurements, the GFP-positive area was measured after threshold-based segmentation using Fiji/ImageJ.

For parasite cluster analysis, vacuoles containing a single GFP-positive parasite were classified as non-clustered, whereas vacuoles containing more than three GFP-positive parasites were classified as clustered or replicating.

CERT and VAP-B recruitment to the PVM was assessed manually from fluorescence microscopy images based on the apparent enrichment of the respective signals along the parasitophorous vacuole boundary. Vacuoles were scored as PVM-positive when a discernible rim-like enrichment was observed.

##### In vivo *Plasmodium* HPA-12 intervention

All animal experiments complied with institutional regulatory standards. Six- to eight-week-old C57BL/6 mice (*n* = 8, gender-balanced) were infected on day 0 by mosquito bite transmission using female *A. stephensi* mosquitoes harboring mature 20-day-old *P. berghei* infections, with 20 infected mosquitoes per mouse. On day 1, mice were randomized into two gender- and weight-matched groups (*n* = 4 per group). Mice received a single tail-vein intravenous administration of either 4 mg/kg HPA-12 (TCI, Cat. #H1553) or an equivalent volume of vehicle control. Animals were monitored daily for 21 days, with a humane endpoint defined as blood-stage parasitemia reaching 40%.

##### Sphingolipid metabolic labeling and click chemistry

For direct ceramide tracking, host cells were pulsed with 5 μM BSA-complexed BODIPY TR-ceramide (Invitrogen). For post-infection pulse experiments, the probe was added during the final 3 h of infection. For pre-labeling experiments, cells were pulsed for 1.5 or 3 h, washed three times with PBS, and chased in fresh medium for 3 h before parasite inoculation.

For metabolic tracking of sphingosine-derived sphingolipids, cells were pulsed with 5 μM alkyne-sphingosine (Cayman Chemical) for 1.5 h, washed three times with PBS, and chased for 3 h before infection. Where indicated, HPA-12 was maintained at 20 μM from the onset of lipid labeling through fixation.

Alkyne-labeled lipids were detected by copper-catalyzed azide-alkyne cycloaddition using the Click-iT Plus Alexa Fluor Picolyl Azide Toolkit (Thermo Fisher Scientific). Fixed and permeabilized cells were incubated for 30 min at room temperature in the dark with a freshly prepared reaction mixture containing Alexa Fluor 647-conjugated picolyl azide, CuSO₄, copper-protecting ligand, and reducing agent. Cells were washed thoroughly with PBS before mounting and imaging.

##### Image segmentation and sphingolipid enrichment quantification

Confocal images were acquired using an Olympus FV3000 microscope with identical laser power and gain settings across experimental cohorts. Spatial segmentation and quantitative analysis were performed using a custom Fiji/ImageJ macro. Host cells containing a single parasite vacuole were selected for analysis. The parasitophorous vacuole mask was defined from the parasite GFP channel using Gaussian filtering and Otsu thresholding followed by morphological processing. The vacuolar boundary was smoothed to define the PVM contour, and a 1-μm outer boundary region surrounding the PVM was used as the region of interest. Integrated sphingolipid fluorescence within this region was normalized to total cellular sphingolipid fluorescence for each host cell.

##### VAP-B proximity labeling and quantitative proteomics

HeLa cells were transfected with miniTurbo-VAPB or miniTurbo-NES plasmids for 24 h and infected with *T. gondii* at an MOI of 0.5 for 3 h following a synchronized PBS wash. At 4 h post-infection, proximity labeling was initiated by incubation with 50 μM biotin for 1 h. Cells were lysed in RIPA buffer supplemented with protease inhibitors, and total protein concentrations were determined using a BCA assay (Pierce). Biotinylated proteins from 3 mg total protein per sample were captured using streptavidin magnetic beads, eluted, trapped in SDS-PAGE gels, and subjected to in-gel trypsin digestion following Coomassie staining.

Peptides were analyzed by LC-MS/MS using an Orbitrap Eclipse Tribrid mass spectrometer. Raw files were processed using Proteome Discoverer 3.2 (Thermo Fisher Scientific) with SequestHT and Amanda 3.0 algorithms and filtered for high/medium FDR confidence, requiring >3 unique peptides and >5% sequence coverage. MS1 area-under-the-curve abundances were log₂ transformed and normalized to propionyl-CoA carboxylase alpha (PCCA)^29^. Missing values were imputed using 0.5 × the limit of detection for left-censored missing data or group averages for random missing values within biological cohorts. Relative enrichment was expressed as the log₂ fold change of miniTurbo-VAPB relative to miniTurbo-NES. Candidate interactors were prioritized based on FFAT motif scores calculated using the method described by Slee and Levine [19], with a score ≤2 used as the high-confidence cutoff.

##### Reverse co-immunoprecipitation

To validate the association between TgVPS13 and host VAP-B, HeLa cells were infected with *T. gondii* VPS13-AID-3×HA parasites and harvested at 5 h post-infection. Monolayers were washed at 1 h post-infection to remove unattached parasites and lysed in ice-cold buffer containing 20 mM Tris-HCl (pH 7.4), 50 mM NaCl, 1 mM EGTA, and 1% Triton X-100 supplemented with protease and phosphatase inhibitors.

Cleared lysates were incubated overnight at 4°C with anti-HA antibody (clone 12CA5, Roche) or mouse IgG control. Immune complexes were captured using Protein A/G Sepharose beads (Santa Cruz Biotechnology), washed, and eluted by boiling in 4X SDS-PAGE sample buffer. Eluted proteins were analyzed by western blotting for endogenous VAP-B, with GAPDH assessed as a cellular protein control.

##### Auxin-inducible TgVPS13 depletion and immunofluorescence

HFF cultures were infected with either the parental TIR1-expressing *T. gondii* line or the VPS13-AID-3×HA strain. Conditional depletion of TgVPS13 was induced using indole-3-acetic acid (IAA; Sigma-Aldrich) at a final concentration of 500 μM. Control parasites were treated with an equivalent volume of ethanol vehicle. Parasites were maintained under the respective treatment conditions for 24 h before host-cell egress. Egressed parasites were subsequently used for infection assays to assess TgVPS13 localization and sphingosine-derived sphingolipid enrichment. Auxin (IAA) or vehicle was maintained during the downstream infection period where indicated. For immunofluorescence analysis, antibodies against HA, VAP-B, and SAG1 were used as appropriate.

##### Reinfection plaque assay and parasite fitness quantification

To assess parasite fitness following disruption of host lipid trafficking, a two-stage reinfection plaque assay was performed across three independent biological replicates. HeLa cells comprising Cas9 control, *VAPB* knockout, shRNA control, *CERT1* knockdown, and *VAPA* knockdown lines, as well as cells continuously treated with 20 μM HPA-12, were seeded in 24-well plates and infected with *T. gondii* at an MOI of 0.5 in duplicate. Monolayers were washed at 3 h post-infection and incubated until 60 h post-infection. Host cells were mechanically lysed, and released tachyzoites were purified by passage through a 27G needle followed by filtration through a 5-μm polycarbonate membrane.

A 1/10 volume of the recovered parasite suspension from each primary well was transferred onto confluent HFF monolayers in secondary 24-well plates. Following 4 days of incubation, monolayers were washed twice with PBS, fixed with 4% PFA for 20 min at room temperature, and stained with 0.5% crystal violet (HiMedia Laboratories) for 10 min. Plates were washed with deionised water and air-dried. Plaque areas were quantified from digital images using Fiji/ImageJ. Images were converted to 8-bit grayscale, and a consistent region of interest was applied to each well to exclude peripheral artifacts. Plaques were identified by thresholding and quantified using the Analyze Particles function. Plaque area was averaged across technical replicates and normalized to the corresponding control condition. Statistical significance was assessed using a one-sample *t*-test against the normalized control value of 1.

##### Docking and hydropathy simulations

The N-terminal Major Sperm Protein (MSP) domain of VAP-B (residues 1-125, sequence mapped from PDB ID: 3IKK) was modeled in complex with candidate FFAT-motif-containing peptides to investigate VAP-B-motif interactions. Structural fragments were restricted to the VAP-B MSP domain and candidate FFAT-containing peptides, excluding disordered, coiled-coil, and transmembrane regions not required for peptide binding. Heteromeric complexes were predicted using AlphaFold-Multimer implemented through ColabFold v1.5.5. Five structural models were generated for each candidate using distinct seeds with joint multiple sequence alignments and structural templates. Models were ranked using the combined confidence metric (0.8 × ipTM + 0.2 × pTM), and complexes with scores ≥0.70 were considered high-confidence structures. Rank-1 models were used for subsequent structural and interface analyses.

Local model confidence was assessed using predicted local distance difference test (pLDDT) scores. Candidate motif interactions were considered high confidence when the mean motif pLDDT was ≥70. The structural integrity of the VAP-B MSP domain was assessed using a modified core pLDDT after excluding flexible residues with pLDDT <70. Predicted aligned error (PAE), interface predicted TM-score (ipTM), and predicted TM-score (pTM) were examined to assess inter-chain positioning and overall model confidence.

Protein-peptide interfaces were characterized using PDBePISA v1.52 to determine interface solvent-accessible surface area, interacting residues, hydrogen bonds, and salt bridges between VAP-B and candidate peptides. Structural visualization and figure preparation were performed using PyMOL v3.2.0a. Hydropathy profiles were mapped onto the protein surface using the Kyte-Doolittle hydropathy scale to evaluate the surface properties of the predicted binding interface.

##### Quantification and statistical analysis

Statistical analyses were performed using GraphPad Prism 10, Python (Library: scipy.stats) and R program. Continuous data are presented as mean ± standard deviation (SD) or standard error of the mean (SEM) from at least three independent biological experiments, as indicated in the figure legends. Differences between two experimental groups were evaluated using two-tailed Welch’s t-tests. Plaque-area and flow cytometry data normalized to the corresponding control were evaluated using a one-sample t-test against normalized value of control (1 or 100). Categorical and phenotypic distribution data (e.g., CERT recruitment categories and vacuole size profiles) were evaluated using Pearson’s Chi-square tests of independence. Host survival curves and time-to-patency were analyzed using log-rank (Mantel-Cox) tests. Statistical significance was defined as *p < 0.05, **p < 0.01, and ***p < 0.001; p > 0.05 was considered not significant (ns). Exact n values, specific statistical tests, and definitions of error bars for individual experiments are provided in the corresponding figure legends.

## Data availability

All data generated and analysed in this study are included in the Article and Supplementary Information. Additional information is available from the corresponding author upon request.

## Acknowledgements

This work was supported by the Science and Engineering Research Board-Anusandhan National Research Foundation (SERB-ANRF) (SRG/2023/001874) to K.V., the Department of Health Research-Indian Council of Medical Research (DHR-ICMR) (2022-1021) to K.V., the DBT-Wellcome Trust India Alliance Intermediate Fellowship (IA/I/23/2/506998) to K.V., and intramural funding from the Indian Institute of Science Education and Research Thiruvananthapuram (IISER Thiruvananthapuram) to K.V. shRNA used in the study were obtained from the shRNA Resource Center at Division of Biological Sciences, IISc, Bangalore. We thank Flow cytometry, Imaging and other core facilities of IISER Thiruvananthapuram for their support and assistance. We thank Feng Zhang, Alice Ting, David Sibley for the Addgene plasmids lentiCRISPR v2, 3xHA-miniTurbo-NLS_pCDNA3, V5-miniTurbo-NES_pCDNA3, pTUB1:OsTIR1-3FLAG, SAG1:CAT, pLinker-AID-3xHA-DHFR-LoxP respectively.

## Author Information

### Contributions

Conceptualization, V.M. and K.V.; methodology, V.M., I.P., V.N., A.P.M., P. B., S.P., U.K., B.B. K.A.K., and K.V.; writing, V.M., P.B., and K.V.; bioinformatic analysis, V.M., P.B., M.K. and K.V.; formal analysis, V.M., P.B., and K.V.; investigation, V.M., and K.V.; resources, K.V.; visualization, V.M., and K.V.; supervision, K.A.K., and K.V.; and funding acquisition: K.V.

### Corresponding author

Correspondence to Kamalakannan Vijayan.

### Ethics Declaration Competing interests

The authors declare no competing interests.

### Ethics

The Institutional Biosafety Committee of the Indian Institute of Science Education and Research Thiruvananthapuram (IISER TVM) and the Review Committee on Genetic Manipulation (RCGM), Department of Biotechnology, Government of India, have approved the use of laboratory research protocols (IBSC/August-05-2022 and IBKP UAC No: INDRDMAL0236) involving the use of *Toxoplasma gondii* and *Plasmodium* species. The procedures involving animals were approved by the Institutional Animal Ethics Committee of IISER TVM (IISERTVM/IAEC/2025-XII/FormB-06). No human subjects, human tissue, or human clinical data were used in this study.

---

**Figure S1.**
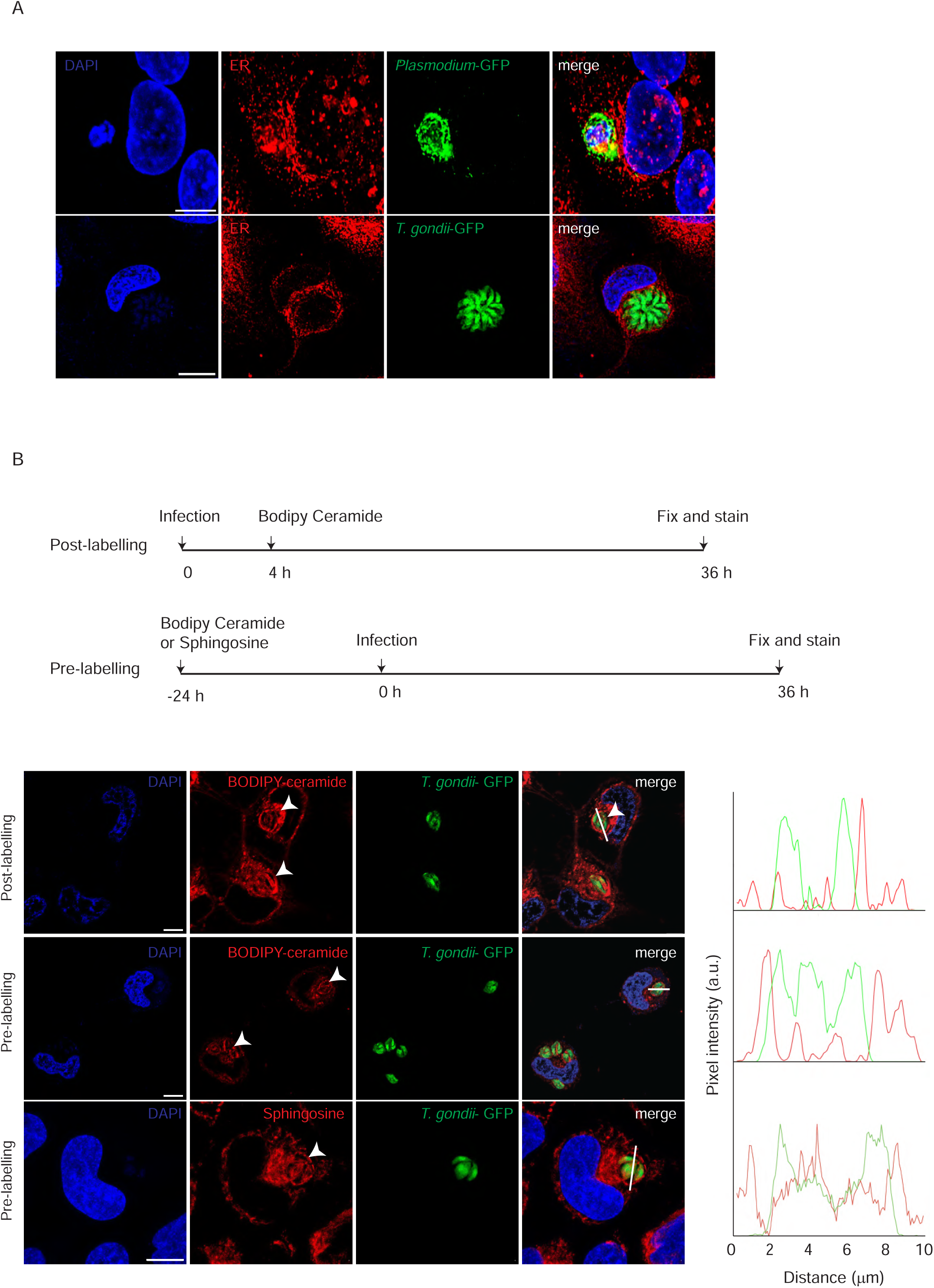
Host ER closely associates with the apicomplexan vacuole, and host-derived sphingolipids accumulate at the PVM. **(A)** Representative confocal micrographs showing host endoplasmic reticulum (ER) networks surrounding the parasitophorous vacuole in *Toxoplasma gondii*- and liver-stage *P. yoelii*-infected host cells. **(B)** Fluorescence microscopy and spatial line-scan analysis of host-derived sphingolipid accumulation at the *T. gondii* parasitophorous vacuole membrane (PVM). Top, BODIPY-ceramide added at 4 h post-infection (hpi) and imaged at 36 hpi. Middle, host cells pre-labeled with BODIPY-ceramide for 24 h and thoroughly washed before parasite infection. Bottom, host cells pre-labeled with fluorescent sphingosine before infection. Arrowheads indicate enrichment at the vacuolar boundary. Scale bars, 5 μm.

**Figure S2.**
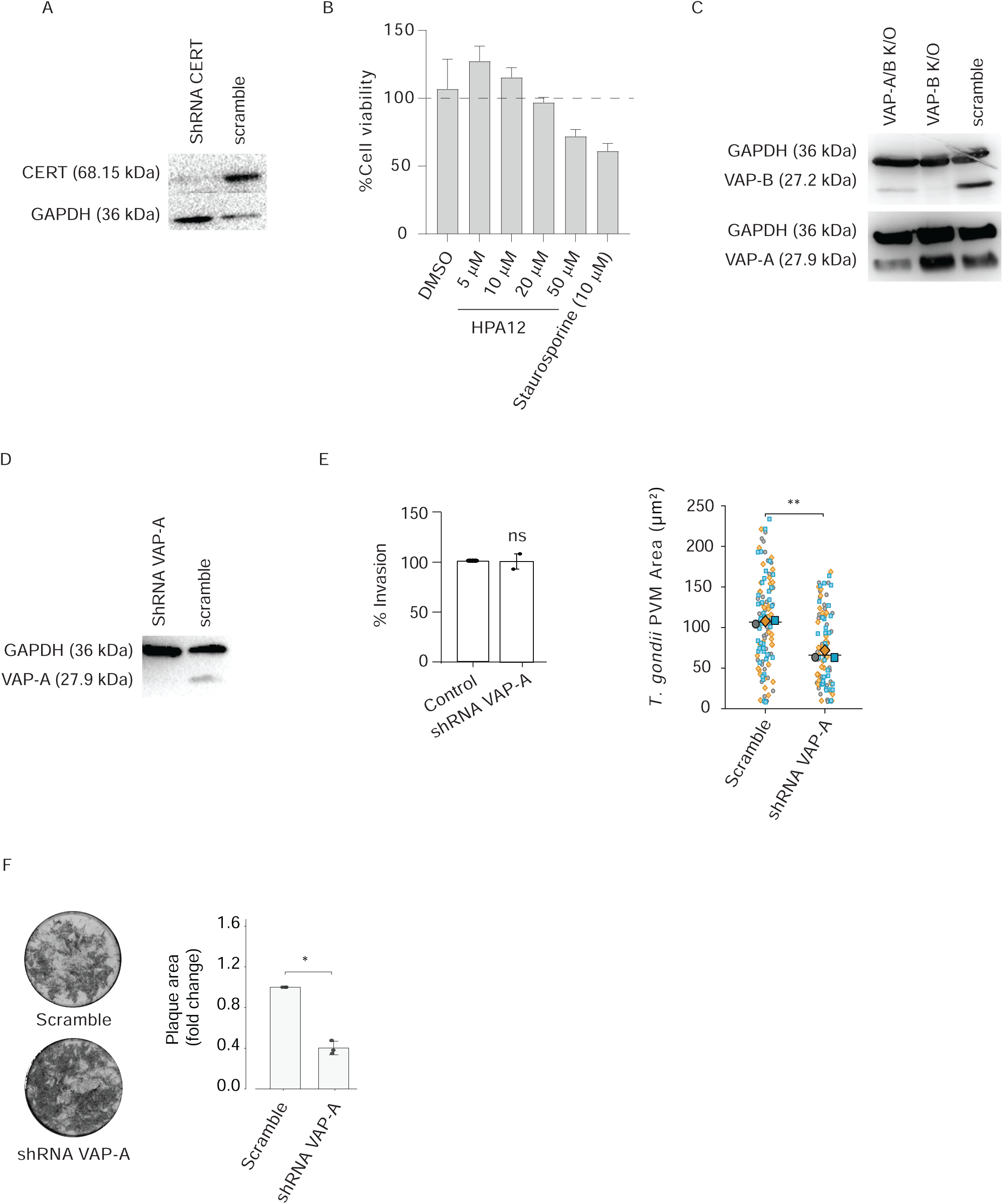
Validation of host CERT and VAP knockdown and knockout cell lines. **(A)** Immunoblot analysis confirming shRNA-mediated depletion of host CERT (68.15 kDa) compared to scramble control host cells. Host GAPDH (36 kDa) serves as a loading control. **(B)** MTT assay to quantify cell viability following HPA-12 treatment. **(C)** Immunoblot verification of host single VAP-B knockout (VAP-B K/O) and double VAP-A/B knockout (VAP-A/B K/O) lines generated by CRISPR-Cas9 compared to scramble controls. Blots are probed for endogenous VAP-B (27.2 kDa), VAP-A (27.9 kDa), and host GAPDH (36 kDa). **(D)** Immunoblot analysis validating shRNA-mediated knockdown of host VAP-A relative to scramble control. Host GAPDH (36 kDa) serves as a loading control. **(E)** Invasion efficiency (left) and the quantification of *T. gondii* parasitophorous vacuole membrane (PVM) area (µm²) (right) at 48 h post-infection in scramble control vs. shRNA VAP-A host cells. Statistical significance was determined using a two-tailed one-sample (invasion) or Welch’s *t-*test (development) (**** *p* < 0.0001). **(F)** Representative crystal violet plaque assay wells (left) and quantification of relative plaque area fold change (right) following *T. gondii* growth in scramble control vs. shRNA VAP-A host cells. Statistical significance was evaluated using a one-sample t-test (* *p* < 0.05). Data represent mean ± SD or mean ± SEM (**E,** right) from *n* = 3 independent experiments.

**Figure S3.**
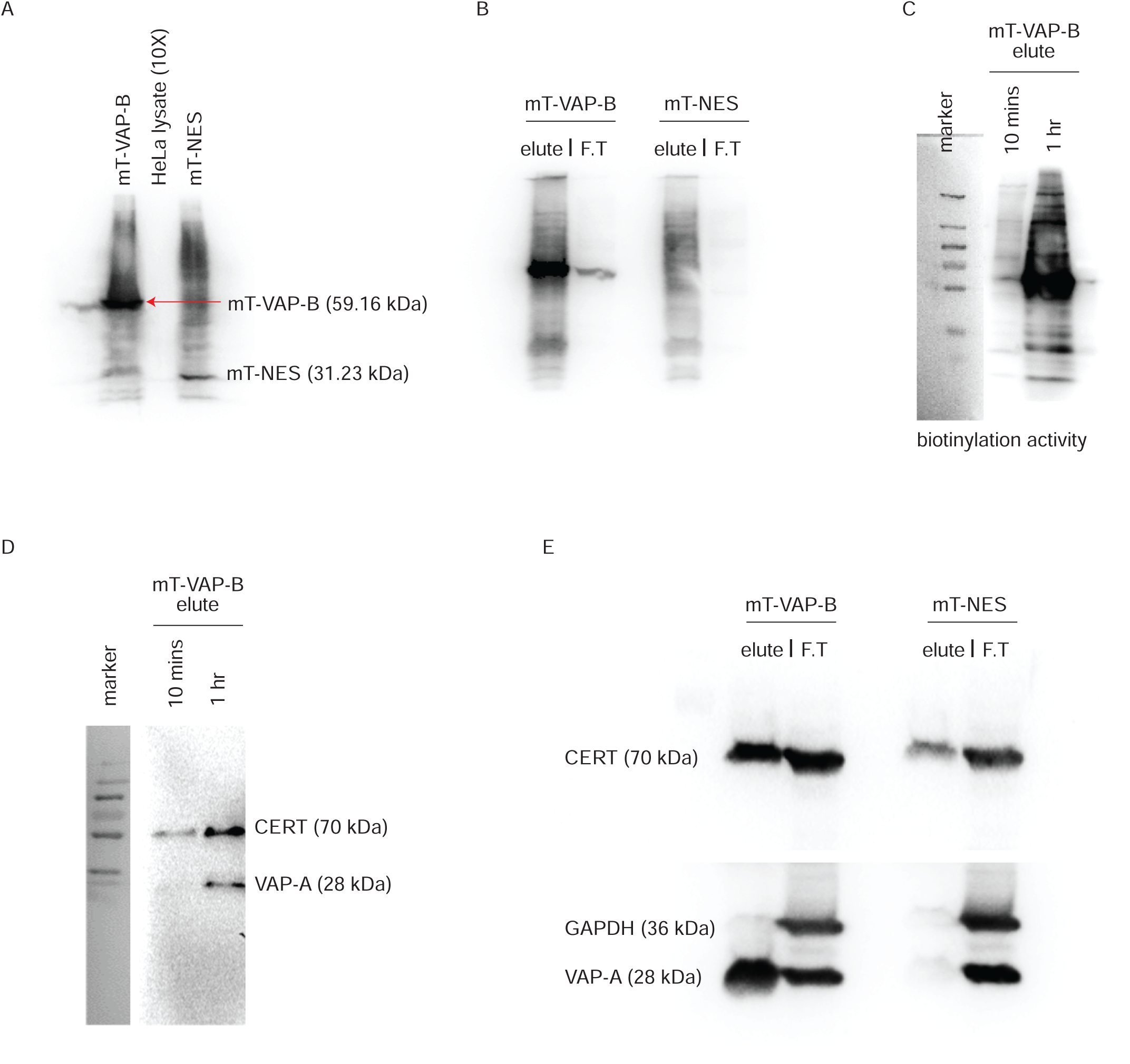
Validation and quality control of the miniTurbo VAP-B proximity labeling system. **(A)** Immunoblot demonstrating expression and streptavidin-bead elution of miniTurbo-tagged VAP-B (mT-VAP-B, 59.16 kDa) and control cytosolic miniTurbo-NES (mT-NES, 31.23 kDa) from host cell lysates across labeling durations (10 min, 1 hr). **(B)** Immunoblot validating specific co-purification of canonical interactor endogenous host CERT (68.15 kDa) in mT-VAP-B eluates compared to mT-NES control eluates and flow-through (F.T.) fractions. **(C)** Biotinylation activity assay showing global biotinylated protein capture in host lysates following 10 min and 1 hr labeling periods. **(D)** Optimization of biotinylation labeling duration (10 min vs. 1 hr). Streptavidin eluates demonstrate that 1 hr biotinylation labeling yields superior capture and enrichment of canonical VAP-B interactors, host CERT (68.15 kDa) and VAP-A (28 kDa). **(E)** Specificity evaluation of mT-VAP-B proximity labeling. Known interactors host CERT (68.15 kDa) and VAP-A (28 kDa) specifically co-purify in mT-VAP-B eluates but are absent in mT-NES control eluates, whereas non-interacting host GAPDH (36 kDa) is cleanly excluded from both eluates and retained in the flow-through (F.T.) fractions.

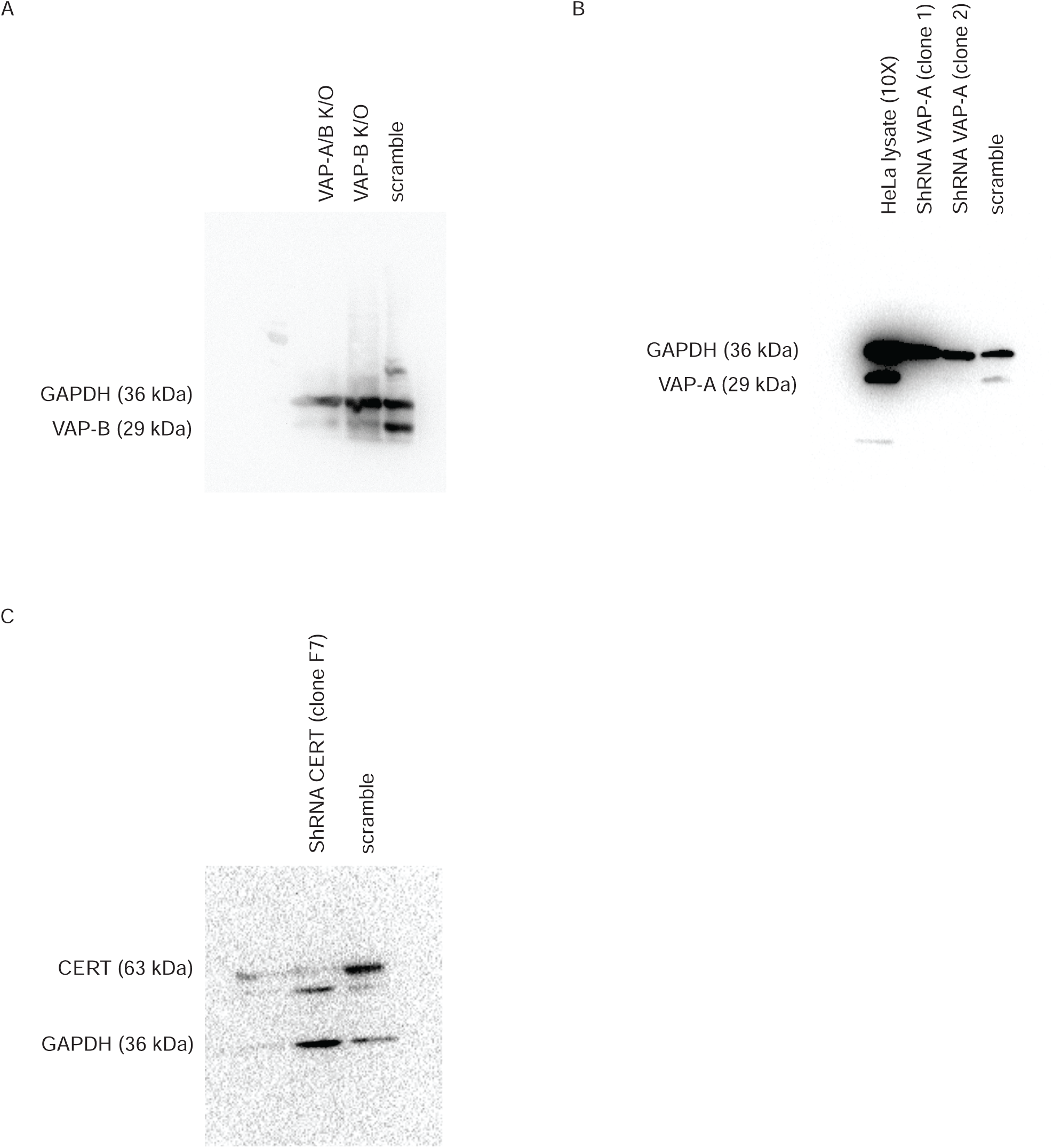

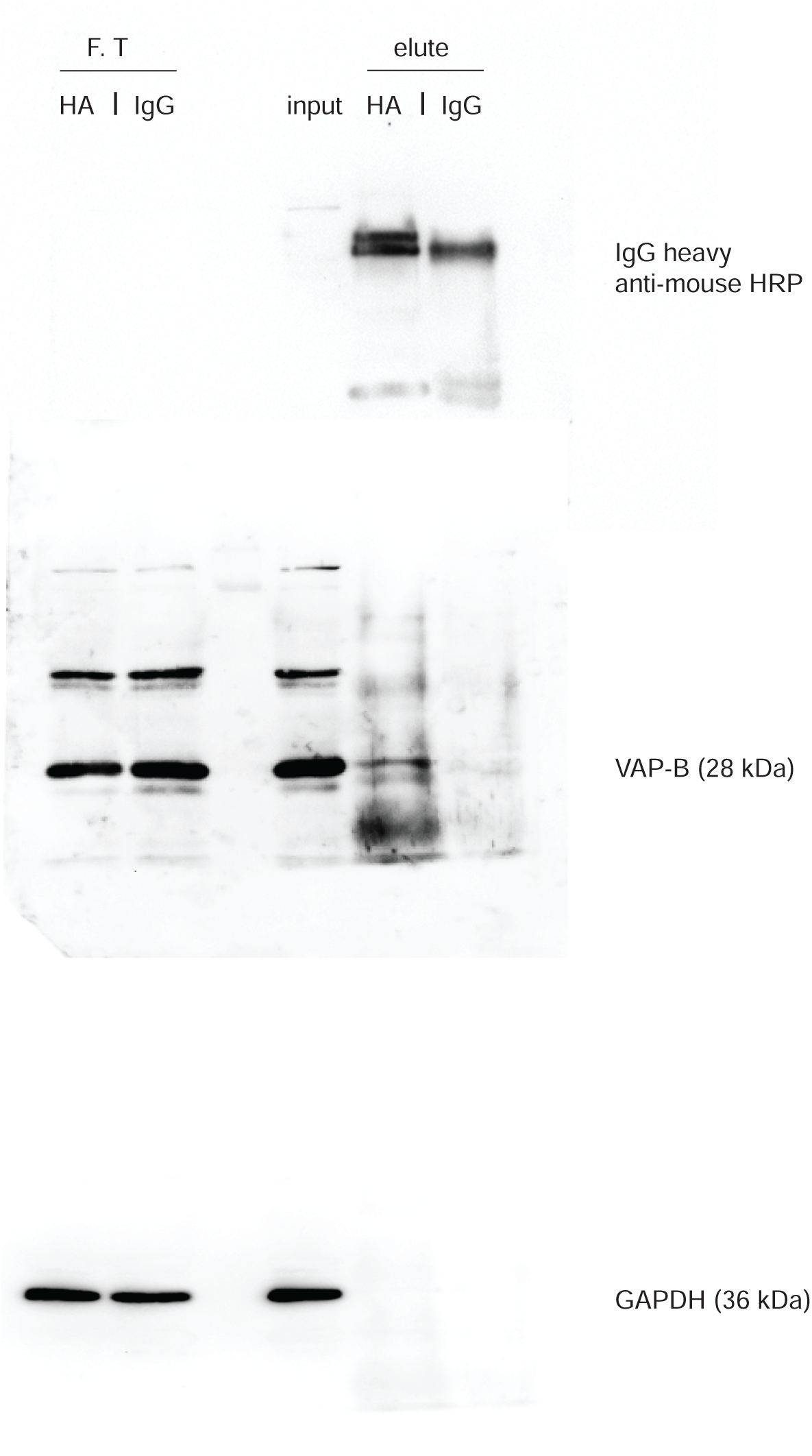

